# Integrative analysis of DNA replication origins and ORC/MCM binding sites in human cells reveals a lack of overlap

**DOI:** 10.1101/2023.07.25.550556

**Authors:** Mengxue Tian, Zhenjia Wang, Zhangli Su, Etsuko Shibata, Yoshiyuki Shibata, Anindya Dutta, Chongzhi Zang

**Author notes:** Correspondence should be addressed to (A.D.) or (C.Z.).

## Abstract

Based on experimentally determined average inter-origin distances of ∼100 kb, DNA replication initiates from ∼50,000 origins on human chromosomes in each cell cycle. The origins are believed to be specified by binding of factors like the Origin Recognition Complex (ORC) or CTCF or other features like G-quadruplexes. We have performed an integrative analysis of 113 genome-wide human origin profiles (from five different techniques) and 5 ORC-binding profiles to critically evaluate whether the most reproducible origins are specified by these features. Out of ∼7.5 million union origins identified by all datasets, only 0.27% were reproducibly obtained in at least 20 independent SNS-seq datasets and contained in initiation zones identified by each of three other techniques (20,250 shared origins), suggesting extensive variability in origin usage and identification. 21% of the shared origins overlap with transcriptional promoters, posing a conundrum. Although the shared origins overlap more than union origins with constitutive CTCF binding sites, G-quadruplex sites and activating histone marks, these overlaps are comparable or less than that of known Transcription Start Sites, so that these features could be enriched in origins because of the overlap of origins with epigenetically open, promoter-like sequences. Only 6.4% of the 20,250 shared origins were within 1 kb from any of the ∼13,000 reproducible ORC binding sites in human cancer cells, and only 4.5% were within 1 kb of the ∼11,000 union MCM2-7 binding sites in contrast to the nearly 100% overlap in the two comparisons in the yeast, *S. cerevisiae*. Thus, in human cancer cell lines, replication origins appear to be specified by highly variable stochastic events dependent on the high epigenetic accessibility around promoters, without extensive overlap between the most reproducible origins and currently known ORC- or MCM-binding sites.

## Introduction

DNA replication is essential for the duplication of a cell and the maintenance of the eukaryotic genome. To replicate the human diploid genome of ∼3 billion base pairs, efficient cellular programs are coordinated to ensure genetic information is accurately copied. In each human cell cycle, replication starts from ∼50,000 genomic locations called replication origins (Hu and Stillman 2023). At an origin of replication, the double-stranded DNA is unwound, and primase-DNA polymerase alpha lays down the first RNA primer extended as DNA. Once replication is initiated, the helicase involved in unwinding the origin continues to unwind DNA on either side and the replication proteins, including DNA polymerases, help copy the single-stranded DNA. Unlike yeast origins, human and most eukaryotic origins do not seem to have a clear DNA sequence preference(Hu and Stillman 2023), (Leonard and Méchali 2013). Yet it becomes clear that chromatin context and other DNA-based activity are important factors for human and non-yeast eukaryotic origin selection. A great amount of effort has been spent to understand how human origins are specified and sometimes different trends are observed regarding genomic features enriched at human origins. For example, G-quadruplex motifs were found to be associated with human origins(Besnard et al. 2012), but can only explain a small subset of origins even after controlling for technical biases(Foulk et al. 2015). To better resolve these issues, we reasoned that a systematic analysis of all available human origin-mapping datasets will help uncover potential technical and biological details that affect origin detection.

To profile replication origins genome-wide, several sequencing-based methods have been developed, including Short Nascent Strand (SNS)-seq (Foulk et al. 2015), Repli-seq (Hansen et al. 2010), Rerep-seq (Menzel et al. 2020), and Bubble-seq (Mesner et al. 2013). SNS-seq assays across different human cell types identified a subset of origins that can explain ∼80% of origins in any tested cell type(Akerman et al. 2020), however no study has systematically examined the consistency within and between different origin-mapping techniques. These methods are based on different molecular capture strategies and therefore are expected to have technical biases. For instance, SNS-seq utilizes lambda exonuclease (λ-exo) to enrich for RNA-primed newly replicated DNA by removing parental DNA fragments. However, the products may be contaminated with chromosomal DNA fragments or be affected by the cutting bias of λ-exo against GC-rich sequences (Foulk et al. 2015), although it has been proposed that one can experimentally correct and control for such biases(Akerman et al. 2020). SNS-seq is the only method that yields origins at a resolution of a few hundred base-pairs, with all other methods delineating larger initiation zones (IZ) that may have multiple origins. Repli-seq relies on nucleotide pulse-labeling and antibody enrichment of newly replicated DNA superimposed on cells fractionated at different parts of S phase by flow cytometry. Here the results may be contaminated by DNA non-specifically associated with the antibody(Zhao et al. 2020). Rerep-seq only captures the new synthesized sequences that replicate more than once when replication is dysregulated, and these origins tend to be enriched in the early replicating, epigenetically open parts of the genome (Menzel et al. 2020). Bubble-seq (Mesner et al. 2013) generates long reads because it captures the origin-containing replication intermediates by selection of DNA bubbles, and so may enrich for origins that are flanked by pause sites. Okazaki-seq (OK-seq) identifies sites where the direction of the Okazaki fragments changes from left-ward to right-ward on the chromosome revealing sites where the two lagging strands diverge from each other (Petryk et al. 2016). Because the technical biases mentioned above are associated with specific techniques, we suspect that origins captured across different techniques are less likely to be affected by those biases. Given the large number of datasets in the public domain (>100 datasets for human origins), this is an opportune time to study the reproducibility between the studies and determine the most reproducible and consistent origins identified by SNS-seq and confirmed by each of the other methods. This most reproducible group of origins (shared origins) from different research groups using different techniques and different cell lines, will be far fewer than all the origins reported to date and minimize complications from stochastic (noisy) firing of origins, but they are best expected to fit the current hypotheses regarding origin specification.

Yeast origin recognition complex (ORC) binds to double-stranded DNA with sequence specificity, helps to load minichromosome maintenance protein complex (MCM) and thus prepare the origins for subsequent firing (Bell and Dutta 2002; Remus et al. 2009; Bell and Stillman 1992), (Costa and Diffley 2022). Consistent with this, a role of ORC in loading MCM proteins was also described in Xenopus egg extracts (Rowles et al. 1999; Coleman et al. 1996). All of this leads to the expectation that in human cells there will be significant concordance between ORC binding sites and efficient origins of replication. Efforts to define double-stranded DNA sequences bound specifically by human ORC find very little sequence specificity (Vashee et al. 2003; Hoshina et al. 2013), and this may be responsible for the lack of sequence specificity in human origins. This difference between yeast and human ORC is attributed to sequence features in ORC4, one of the subunits of the origin recognition complex (Lee et al. 2021). When a yeast-specific 19 amino acid insert was removed from yeast ORC4 to make it more like the human ORC4, the sequence specificity of ORC binding was lost even in yeast. Despite this loss of sequence specificity, the mutant yeast ORC loaded MCM proteins, and the yeast replicated DNA and survived. Thus, even if human origins of replication do not have specific sequences, one should expect some concordance between ORC binding sites and the most reproducible origins of replication. An additional complexity is that genome-wide analysis found active origins and dormant origins determined by SNS-seq and OK-seq have little difference in ORC or MCM density (Sugimoto et al. 2018; Kirstein et al. 2021). Finally, a few publications have reported significant MCM2-7 loading and DNA synthesis after genetic mutation of Drosophila *ORC1* and human *ORC1*, *ORC2* or *ORC5* genes that reduce the expressed proteins to near undetectable levels (Park and Asano 2008; Shibata et al. 2016; Okano-Uchida et al. 2018; Shibata and Dutta 2020). In two of the instances, the DNA replication seen was in the context of endo-reduplication, a process that is still believed to require the loading of MCM2-7 and the activation of the same into an active CMG helicase. It should be noted, however, that sgRNA screens revealed that the *ORC2* gene was essential for viability in the cancer cell lines mutated for *ORC2*, suggesting that vanishingly small amounts of the *ORC2* gene-product is required for some process essential for cell proliferation (Chou et al. 2021). Definition of the most reproducible and consistent origins identified by different methods from different research groups in different cell lines thus provides a unique opportunity to determine how much overlap is seen between the highly reproducible human origins and the ORC-binding sites reported in the literature from ChIP-seq studies.

To understand the genome-wide distribution patterns of replication origins in an unbiased way, we performed an integrative analysis of 113 DNA replication origin datasets. Because SNS-seq is unique in identifying origins of a few hundred bases, while the other methods identify larger initiation zones that contain origins, we first prepared a list of high-resolution SNS-seq origins that have been identified in at least 20 of the 66 SNS-seq datasets in this study. We next determined those reproducible SNS-seq origins that overlap with initiation zones identified by each of the three techniques (Repli-seq, OK-seq and Bubble-seq) to identify 20,250 high-confidence origins that are shared between SNS-seq and every other method identifying initiation zones. Using these shared origins, which overlap significantly with transcription promoters, we tested whether G-quadruplex sites, CTCF binding sites, ORC binding sites or MCM binding sites help specify origins.

## Results

### A total of 7,459,709 origins from 113 datasets show similar but different genomic features associated with each origin-mapping technique

We collected 113 publicly available replication origin identification studies in different human cell types from five different techniques (Fig. 1-figure supplement 1a), including SNS-seq, Repli-seq, Rerep-seq, OK-seq and Bubble-seq. The complete list of datasets used in this analysis can be found in Supplementary File 1. We processed all the datasets using the first two steps of a pipeline with different parameters considering the various resolution of different techniques, as SNS-seq is unique in identifying high-resolution origins of replication (Fig. 1a). Each of the 113 datasets yielded at least 1000 origins. We merged origins that overlap for at least 1bp from each other and cut the merged regions into 300bp segments, considering origin lengths were significantly longer for Bubble-seq and OK-seq, methods known to identify initiation zones (Fig. 1-figure supplement 1b). A total of ∼7,460,000 union origins were discovered from all techniques (Fig. 1b).

**Figure 1:**
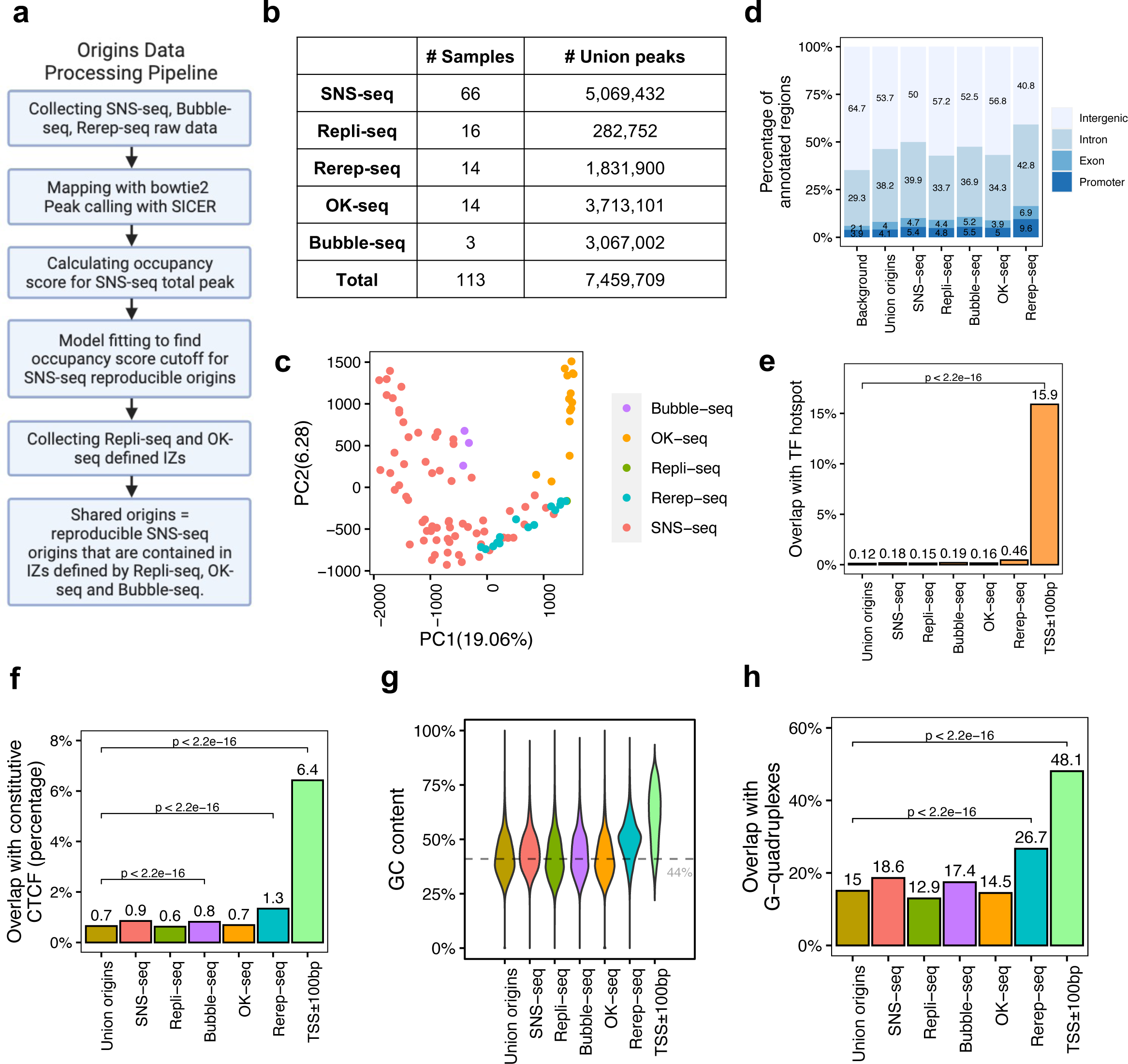
Total 7,459,709 origins defined by four types of techniques show different genomic features. (a) Data processing pipeline. 113 publicly available profiles of origins are processed following the pipeline. (b) Number of samples collected for each technique. In total 7,459,709 union origins were identified. (c) PCA shows the clustering of origin datasets from different techniques. (d) Genomic annotation (TSS, exon, intron and intergenic regions) of different groups of origins. Background is the percentage of each annotation on the whole genome. (e) Overlap with TF hotspots for different groups of origins and promoters. (f) Overlap with constitutive CTCF binding sites for different groups of origins and promoters. (g) GC content of different groups of origins and promoters. Grey line marks the average GC content of human genome. (h) G-quadruplex overlapping rates of different groups of origins and promoters.

Principal Component Analysis (PCA) of all origin datasets shows that origin locations from the same technique are more similar to each other than from different techniques (Fig. 1c). This is confirmed by pair-wise correlations of the datasets, where it is evident that each method identifies origins that are best correlated with origins identified by that method alone (Fig. 1-figure supplement 2) Although the most popular technique, SNS-seq, shows some variability in the PCA (Fig. 1c), it is also the one that is best correlated with origins identified by other SNS-seq datasets. Re-rep-seq and Bubble-seq seemed to identify initiation zones that were also better correlated with SNS-seq origins (Fig. 1-figure supplement 2). Since Re-rep-seq tends to identify origins that are enriched in early replicating, gene-rich parts of the genome, this result suggests that SNS-seq and Bubble-seq also have some bias for origins in early replicating, gene-rich parts of the genome. However, we did not observe clear trends of clustering of origins identified in similar cell types (Fig. 1-figure supplement 1c), even when we focused only on SNS-seq origins (Fig. 1-figure supplement 1d). The one exception was the SNS-seq origins from T lymphoblasts, which were very closely clustered with each other in six datasets, but these were all done in a the CCRF-CEM cell-line (or a derivative) by one group at the same time(Murai et al. 2018). We only examined SNS-seq data from 2018 onwards, when the lambda exonuclease-based digestion step had been incorporated into SNS-seq protocols, but wondered if there was a steady improvement of reproducibility as one progressed through the years. However, SNS-seq datasets do not become progressively more reproducible as one goes year-by-year from 2018-2022 (Fig. 1-figure supplement 1e). The results suggest that there is significant difference in origins captured by different techniques that cannot be explained by differences in cell choices. For the most popular technique (SNS-seq) the variability may be least when confined to a single cell line studied at the same time by the same group.

We next checked the genomic characteristics of origins defined by each technique. Regardless of technique used, around half of the detected origins fall in intergenic regions, followed by 30-40% allocated to introns (Fig. 1d). We recently reported 40,110 genomic regions are routinely bound by more than 3000 TF ChIP-seq data, named as TF binding hotspots (Hu et al. 2021). These TF binding hotspots likely represent highly active open chromatin regions. Despite the differences among techniques, the origins in general have lower overlap with TF binding hotspot (<0.5%), compared to the 16% overlap of gene promoter regions (TSS +/-100bp) with such TF binding hotspots (Fig. 1e). We further focused on one specific TF, CCCTC-binding factor (CTCF), which is important in organizing DNA structure and reported to be associated with replication (Su et al. 2020). Origins show significantly lower overlap with constitutive CTCF sites, defined as those that are conserved across cell types (Fang et al. 2020), compared to promoters (Fig. 1f). G-quadruplexes are found to be correlated with both replication and transcription (Lipps and Rhodes 2009). However, compared to promoters, origins defined by these techniques show significantly lower GC content (Fig. 1g). In addition, there is significantly lower overlap of these origins with G-quadruplex sites identified from predicted G-quadruplex motif regions (Bedrat et al. 2016), compared to the overlap of promoters with G-quadruplex sites (Fig. 1h). Thus the reported correlations between CTCF binding sites or G-quadruplexes with origins are not as striking as that of these features with promoters.

We found that all origins, regardless of technique by which identified, are slightly enriched at gene promoters, exons, or introns, compared with the genome background (Fig. 1d). Among the five different techniques, origins defined by Rerep-seq show the highest level of enrichment with promoters, TF hotspots, constitutive CTCF binding sites and G-quadruplexes and the highest GC content. This is consistent with our knowledge that areas of the genome that are re-replicated when the cell-cycle is disturbed, are enriched in parts of the genome that replicate early in S phase, regions which are enriched in transcriptionally active (and thus epigenetically open) genes and their promoters.

### Shared origins are associated with active chromatin and transcription regulatory elements

To address the potential biases caused by each technique, we investigated how many of the ∼5 million union SNS-seq origins are reproducible in SNS-seq data and confirmed by other sequencing-based techniques. As we will describe below, these shared origins show significant overlap with promoters. Because the Re-rep-seq origins appear to be slightly different from the origins identified by other techniques, with highest overlap with transcriptionally active genes and promoters, we decided to exclude the Re-rep-seq from the analysis of shared origins and still reached the conclusion summarized above.

SNS-seq origins have the highest resolution. We used the following strategy to determine how many independent confirmations of an SNS-seq origin is sufficient for selecting an SNS-seq origin as a reproducible origin. The occupancy score of each origin defined by SNS-seq (Fig. 2-figure supplement 1a) counts the frequency at which a given origin is detected in the datasets under consideration. Plotting the number of union SNS-seq origins with various occupancy scores with all SNS-seq datasets published after 2018, we sought to determine whether the curve deviates from the random background at a given occupancy score (Fig. 2a). For the random background, we assumed that the number of origins confirmed by increasing occupancy scores decreases exponentially (see Methods and Supplementary File 2). The threshold occupancy score, to determine whether an origin is a reproducible origin, is the point where the observed number of origins deviates from the expected background number (with an empirical FDR < 0.1) (Fig. 2a), an occupancy score of 20. Thus the reproducible SNS-seq origins (with a FDR<0.1) were those observed in at least 20 SNS-seq datasets.

**Figure 2:**
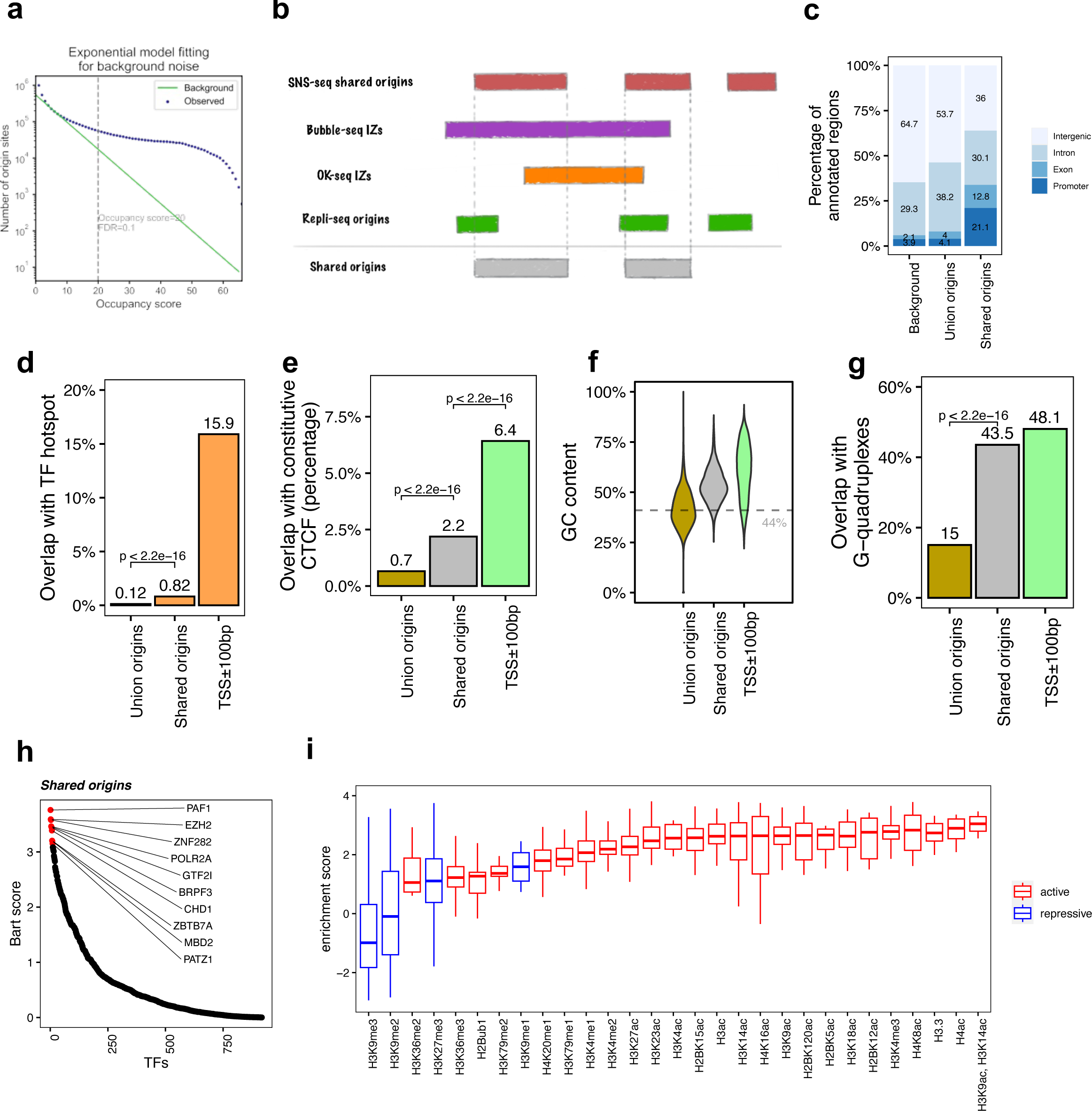
The shared origins are enriched with certain transcription factors and active histone marks. (a) SNS-seq origins fitting distribution to an exponential model shows that an occupancy score ≥20 selected for reproducible SNS-seq origins. (b) Conceptual model of how shared origins is calculated. Any SNS-seq shared origin that overlaps with Bubble-seq IZ, OK-seq IZ and Repli-seq origin together is considered as an origin identified by all four techniques (shared origins). (c) Genomic annotation of union origins and shared origins. (d) Overlap with TF hotspots of union origins and shared origins. (e) Overlap with constitutive CTCF binding sites of union origins and shared origins. (f) GC content of union origins and shared origins. (g) G-quadruplex overlapping rates of union origins and shared origins. (h) BART prediction of TFs associated with shared origins. (i) Enrichment of histone marks at shared origins using all origins as control.

We next determined which reproducible SNS-seq origins are confirmed by origins from Repli-seq and replication initiation zones (IZs) from Bubble-seq and OK-seq (Fig. 2b). 20,250 of the reproducible SNS-seq origins were found to overlap with an origin or IZ identified by each of the three other techniques and were called shared origins. These high-confidence shared origins consist of 0.27% of all ∼7.5 million union origins. The coordinates of the shared origins are available in the supplementary files.

The shared origins have a greater overlap rate with gene promoters and exons compared to union origins (Fig. 2c). This is in line with the previous observation that replication and transcription are highly coordinated and enrichment of origins at TSS (Ganier et al. 2019; Cook 1999; Karnani et al. 2010). Shared origins also have a substantially higher overlap rate than union origins with TF binding hotspots (Fig. 2d). Moreover, shared origins have a higher overlap rate with constitutive CTCF binding sites, compared to union origins (Fig. 2e), and have a higher GC content and overlap with G-quadruplexes than union origins (Fig. 2f-g).

BART (Wang et al. 2018) analyzes the enrichment of TF binding sites (determined experimentally by ChIP-seq) with areas of interest in the genome. We used BART to perform TF binding site enrichment analysis on the shared origins and union origins and identified potential TFs or components of chromatin remodeling factors (e.g., PAF1, EZH2, ZNF282, POLR2A, GTF2I) whose binding sites are associated with the shared origins (Fig. 2h, Fig. 1-figure supplement 1f). The high enrichment of activators or repressors of transcription in the factors that have binding sites near shared origins provides more support that the shared origins have properties similar to transcriptional promoters.

To investigate whether the shared origins have a specific chromatin epigenomic signature compared to union origins, we used 5,711 publicly available ChIP-seq datasets for 29 different histone modifications and generated a comprehensive map of histone modification enrichment at shared origins compared to union origins. A substantial enrichment of activating histone marks, including H3K4me3 and H3/H4 acetylation, was observed at shared origins compared to union origins (Fig. 2i). The enrichment of H3K14ac is interesting given the enrichment in the BART analysis of binding sites of BRPF3, a protein involved in this specific acetylation and reported to stimulate DNA replication(Feng et al. 2016), but this modification was not uniquely enriched at the shared origins. These results show that as we move from all origins to a small set of high-confidence shared origins, we see a progressive increase in enrichment of TSS in epigenetically active parts of the genome.

However, transcriptional and epigenetic activators are not the whole story. H3K27me3, a repressive mark, is also enriched at shared origins, although this enrichment is not as high as that of the activating marks. The enrichment of EZH2 binding sites near shared origins is consistent with this observation because EZH2 is part of the Polycomb Repressive Complex 2 known to be a writer of H3K27me3. The paradoxical enrichment of repressive marks is consistent with the shared origins also being near binding sites of MBD (binds methylated DNA and represses transcription), PATZ1 and ZNF282, proteins known to be repressors of transcription.

Having identified a small group of highly reproducible origins (independent of cell-type and technique) we asked whether in a given cell line was one technique superior to the others in identifying such origins. K562 cells have been interrogated for origins by three different techniques: SNS-seq, OK-seq and Repli-seq (Fig. 2-figure supplement 2). Among the 20,250 high-confidence shared origins, 9,901 (48.9%) overlapped with SNS-seq origins in K562; 3,872 (19.1%) overlapped with OK-seq IZs in K562; 1,163 (5.7%) overlapped with Repli-seq origins in K562. This suggests that in one cell line, even the large initiation zones defined by OK-seq or Repli-seq do not capture most of the high-confidence origins. In the opposite direction, where we estimate what fraction of origins found by a given technique fall in a reproducible origin, the opposite result emerges: 2.7% of SNS-seq origins, 17.2% of OK-seq initiation zones and 7.7% of Repli-seq initiation zones overlapped with the 20,250 shared origins (Fig. 2-figure supplement 2). Thus SNS-seq may be able to identify more of the reproducible origins, but it comes with a high false positive rate.

### Human, but not yeast, high-confidence origins have low overlap with known ORC binding sites

Origin recognition complex (ORC) is expected to bind near replication origins during the cell cycle to help define origins (Bell 2002). To investigate the correlation of ORC binding with shared origins detected across different origin-mapping techniques, we analyzed all five publicly available ORC1 and ORC2 ChIP-seq datasets with at least 1,000 peaks (Supplementary File 1c). We identified a total of 34,894 ORC binding sites in the human genome (Supplementary File 3). Union (all) of ORC binding sites are enriched at promoters, TF hotspots, constitutive CTCF sites, GC content and G-quadruplexes compared to randomized genome background (Fig. 3a-e), somewhat similar to what we observed for shared origins (Figure 2).

**Figure 3:**
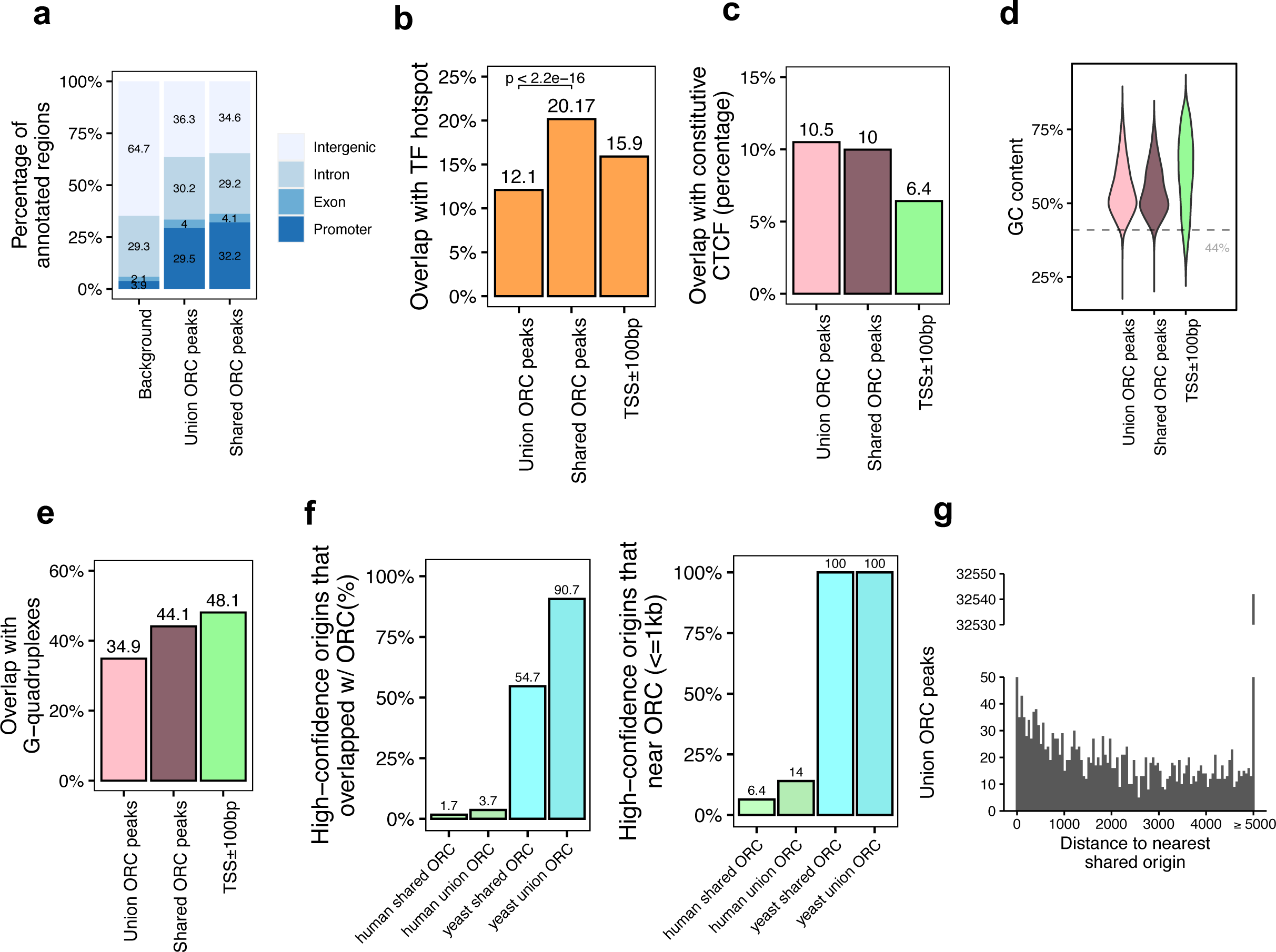
Genomic features of shared ORC binding sites and their co-localization with shared origins. (a) Genomic annotation of union ORC and shared ORC binding sites. (b) Overlap with TF hotspot of union ORC and shared ORC binding sites. (c) Overlap with constitutive CTCF binding sites of union ORC and shared ORC binding sites. (d) GC content of union ORC and shared ORC binding sites. (e) Overlap with G-quadruplex of union ORC and shared ORC binding sites. (f) The percentage of high-confidence origins (shared origins in human and confirmed origins in yeast) that overlapped with (left) or are proximate to (≤1kb) (right) to two types of ORC binding sites (union or shared). (g) Distribution of the distance between ORC binding sites and the nearest shared origin.

Of the 34,894 ORC binding sites in the human genome 12,712 sites were defined as shared ORC binding sites as they occur in at least two ORC datasets (Occupancy score ≥2). For the majority of genomic features, including enrichment at promoter (Fig. 3a), overlap with constitutive CTCF sites (Fig. 3c), GC content (Fig. 3d) and overlap with G-quadruplex (Fig. 3e), ORC binding sites have similar genomic distribution regardless of how many samples they appear in. Interestingly, the G-quadruplex enrichment at either union or shared ORC binding sites is a bit lower than in gene promoter regions (Fig. 3e). The only significant difference between the union ORC sites and the shared ORC binding sites is the significantly higher overlap of the latter with TF binding hotspots (Fig. 3b), suggesting that ORC-ChIP seq data also tend to enrich for highly open chromatin regions.

Based on the assumption that ORC binding sites that appeared in more than one dataset are likely to be true-positive ORC binding sites, we analyzed how many of the 12,712 shared ORC binding sites *overlap* with the 20,250 shared origins identified by our analysis above. The overlap was surprisingly low: 1.7% of the shared origins overlapped with shared ORC binding sites (Fig. 3f, Fig. 3-figure supplement 1a). Even when we relaxed the criteria to look at shared origins that are proximal (≤1 kb) to shared ORC binding sites only 1300 (6.4%) of the shared origins were proximal to the shared ORC binding sites (Fig. 3f) (Supplementary File 4b). The low degree of overlap or proximity of shared origins with shared ORC binding sites indicates that the vast majority of the shared origins are not near experimentally determined ORC binding sites.

In the reverse direction, we asked whether an ORC binding site definitively predicts the presence of an origin nearby. A histogram of the distance between any ORC binding site (union) and the nearest shared origin showed that only 1,086 (3.11%) ORC binding sites are proximal to (≤1 kb) a shared replication origin (Fig. 3g, Supplementary File 4a). Given there are 34,894 ORC binding site and 20,250 shared origins, if ORC binding was sufficient to determine a high confidence origin, nearly half of the ORC binding sites should have been proximate to the shared origins. This low level of proximity between ORC binding and reproducible origins suggests that the current data on ORC binding sites is unable to predict the presence of a high confidence, actively fired origin.

We examined whether the co-localization is better if the analysis is done with data exclusively from the same human cell type (Fig. 3-figure supplement 1b and 1d). Only 8.8% of the 105,881 union origins identified by SNS-seq, OK-seq, or Repli-seq in K562 cells overlap with ORC2 binding sites mapped in the same cell line, and only 4.9% of the 68,003 union SNS-seq origins mapped in HeLa cells overlap with ORC1 binding sites in the same cell line. The overlaps improve marginally if we focus on shared origins: 12.8% of 9,605 shared origins in K562 cells and 6.1% of 3,390 shared origins in HeLa cells overlap with ORC2 and ORC1 ChIP-seq sites in the concordant cell lines.

Experiments in the yeast S. cerevisiae have demonstrated that ORC binding is critical for defining an origin of replication. Indeed, in contrast to humans, in the more compact genome of the yeast, *S. cerevisiae*, there is higher overlap or proximity (within 1 kb) seen between the Yeast ORC ChIP-seq binding sites (Supplementary File 1) and the well mapped yeast origins of replication in OriDB (Nieduszynski et al. 2007). In this case, 100% of the shared origins are proximate to (≤1 kb) all (union) or shared ORC binding sites in yeast (Fig. 3f).

Overall, we found that the shared origins are highly co-localized with ORC binding sites in yeast but not in the human cell lines, suggesting that the current methods of ORC binding site determination are failing to identify functional ORC binding.

### Properties of the 6.4% shared origins co-localized reproduced ORC binding sites

Because of the poor co-localization of shared origins with ORC binding sites, we next focused on the 6.4% of shared origins (1,300) that are proximate to a shared ORC binding site, a group we will call the “highest confidence origins”. Compared with shared origins or ORC-binding sites alone, these 1,300 origins are even more co-localized with gene promoters (Fig. 4a), TF binding hotspots (Fig. 4b) and constitutive CTCF binding sites (Fig. 4c). The GC content is not significantly higher for these “highest confidence origins” (Fig. 4d), and neither was their overlap with G-quadruplexes (Fig. 4e).

**Figure 4:**
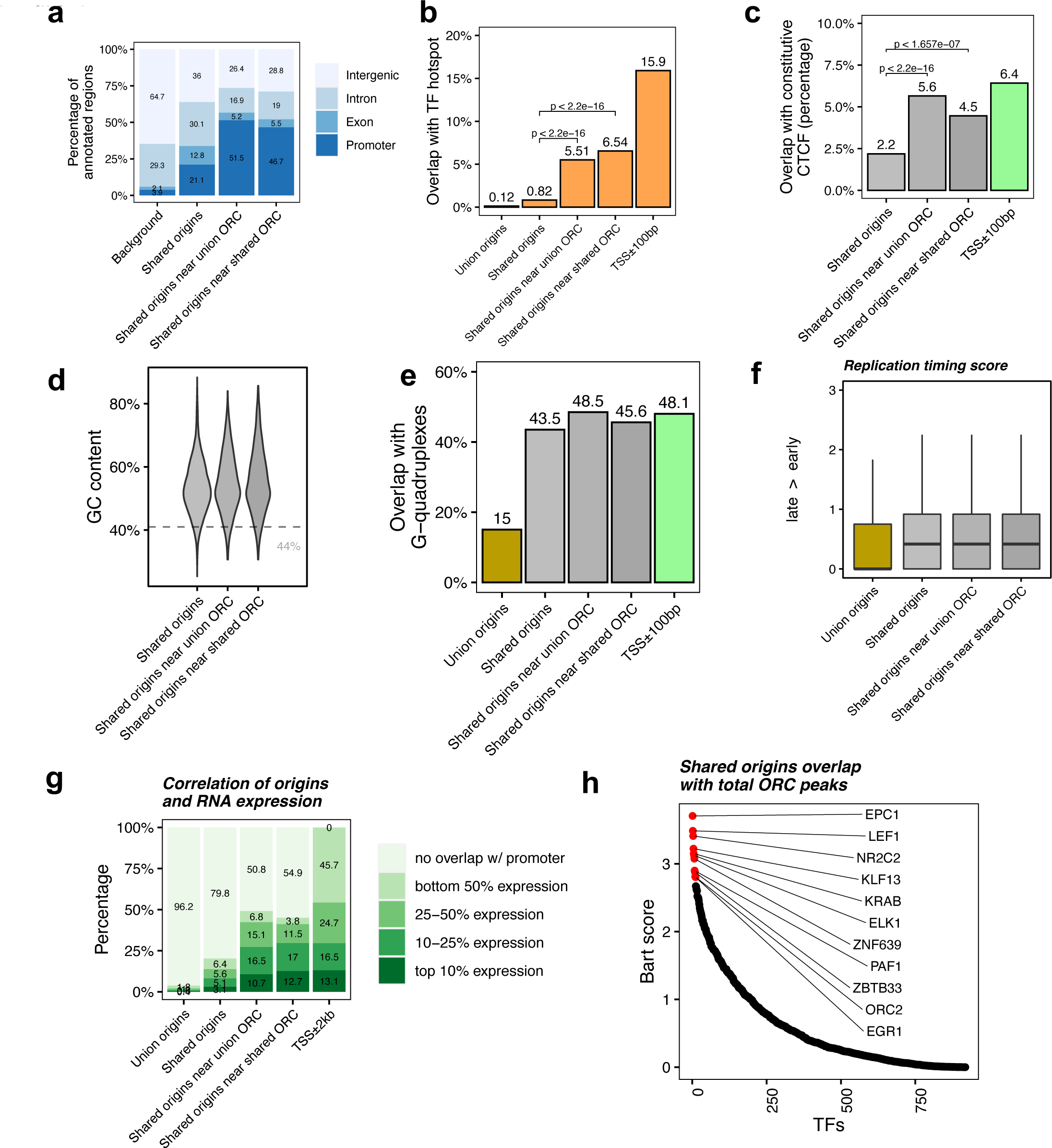
Shared origins near shared ORC binding sites are more correlated with active transcription. (a) Genomic annotation of shared origins and shared origins near (≤1kb) ORC binding sites. (b) Overlap with TF hotspots of shared origins and shared origins near ORC binding sites. (c) Overlap with constitutive CTCF binding sites of shared origins and shared origins near ORC binding sites. (d) GC content of shared origins and shared origins near ORC binding sites. (e) Overlap with G-quadruplex sites of shared origins and shared origins near ORC binding sites. (f) Y-axis: Replication timing score (from Navarro, 2021) for indicated classes of origins. (g) Annotation of expression level of genes that overlapped with different groups of origins. (h) BART prediction of TFs associated with highest confidence origins.

To understand the correlation between the different classes of origins and replication timing, we calculated the replication timing score for the origins following the ENCODE pipeline (Navarro Gonzalez et al. 2021; Hansen et al. 2010), and found that the mean replication timing score of shared origins suggest that they replicate earlier in S phase compared to the union origins, but there is not much difference between shared origins separated by whether they overlapped with ORC binding sites or not (Fig. 4f).

This suggests that even though shared origins near ORC binding sites are more enriched in TSS, TF binding hotspots and CTCF binding sites they are still similar to shared origins globally in their localization in early replicating, epigenetically active parts of the genome.

To investigate if the “highest confidence origins” (shared origins near ORC binding sites) are associated with higher transcriptional activity, we divided all human genes into four groups based on their expression level in the human K562 cell line and found the genes co-localized with the highest confidence origins exhibit higher expression levels compared to genes co-localized with all shared origins or union origins (Fig. 4g).

BART analysis (Wang et al. 2018) of which protein binding sites are most enriched in 743 origins overlapping with any ORC binding sites shows, as expected, that ORC2 binding sites are enriched near these origins. The other proteins bound near these origins are either transcriptional activators like EPC1, LEF1, ELK, EGR1, PAF1 or transcriptional repressors like KLF13, KRAB, ZNF639 or ZBTB33 (Fig. 4h). These results suggest that the shared origins overlapping with ORC binding sites tend to be more associated with transcriptional regulation than all shared origins.

### Overlap of MCM binding sites with shared origins to define another type of highest confidence origins

The six subunit minichromosome maintenance complex (MCM) is loaded on chromatin in G1 and forms the core of the active helicase that unwinds the DNA to initiate DNA replication (Madine et al. 2000). Since MCM2-7 may be loaded by ORC and move away from ORC to initiate DNA replication, it could be expected that even if the ORC binding sites are not proximate to the 20,250 shared origins, they will be proximate with known MCM binding sites. To test this, we analyzed 18 human MCM ChIP datasets (ENCODE Project Consortium 2012; Ivanov et al. 2018; Utani et al. 2017). We identified 11,394 total MCM3-7 binding sites (union) and 3,209 shared MCM binding sites that are defined by an intersection of MCM3, MCM5 and MCM7 union peaks. Overall MCM binding sites displayed very similar genomic features as ORC binding sites (Fig. 5a-e). We then defined the genomic features common among the shared origins that are close to (≤1kb) MCM binding sites. Like ORC-designated origins (Fig. 4b), the MCM-designated origins showed higher overlap with TF binding hotspots (Fig. 5f). Very interestingly, similar to the high overlap between yeast origins and yeast ORC sites (Fig. 3f), around 95% of yeast origins(Lee et al. 2021; Gros et al. 2015) are close to (≤1kb) the union of all yeast MCM sites. In contrast, only ∼4.5% shared human origins are close to (≤1kb) the union of experimentally defined human MCM binding sites (Fig. 5g).

**Figure 5:**
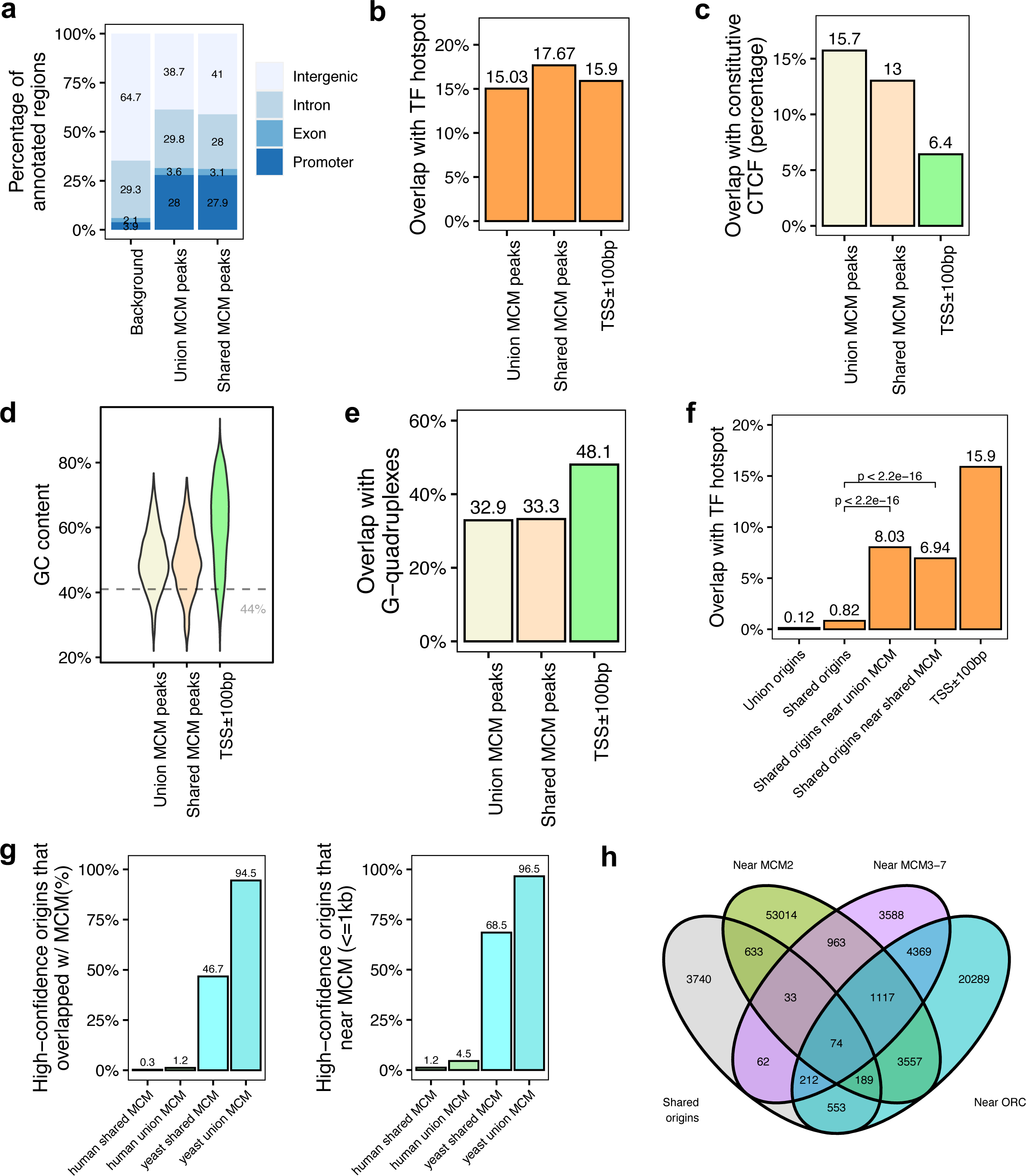
Genomic features of shared MCM binding sites and their co-localization with shared origins. (a) Genomic annotation of union MCM and shared MCM binding sites. (b) Overlap with TF hotspot of union MCM and shared MCM binding sites. (c) Overlap with constitutive CTCF binding rates of union MCM and shared MCM binding sites. (d) GC content of union MCM and shared MCM binding sites. (e) Overlap with G-quadruplex of union MCM and shared MCM binding sites. (f) Overlap with TF hotspots of shared origins and shared origins near MCM binding sites. (g) The percentage of high-confidence origins (shared origins in human and confirmed origins in yeast) that overlapped with (left) or are proximate to (≤1kb) (right) to two types of MCM binding sites (union or shared). (h) Venn diagram of shared origins that are near ORC, MCM2 or MCM3-7 binding sites.

We examined all three types of high confidence origins (ORC designated, MCM3-7 designated and MCM2 designated) in a Venn Diagram (Fig. 5h) and identified 74 that were reproducibly identified by multiple methods (shared origins, near shared ORC binding sites, overlapping with MCM3-7 binding sites and MCM2 binding sites). The coordinates of these origins and their supporting data (ORC and MCM binding sites) are listed in Supplementary File 5, and genome browser views for three of them are shown in Fig. 6 to indicate the relationship between the coordinates.

**Figure 6:**
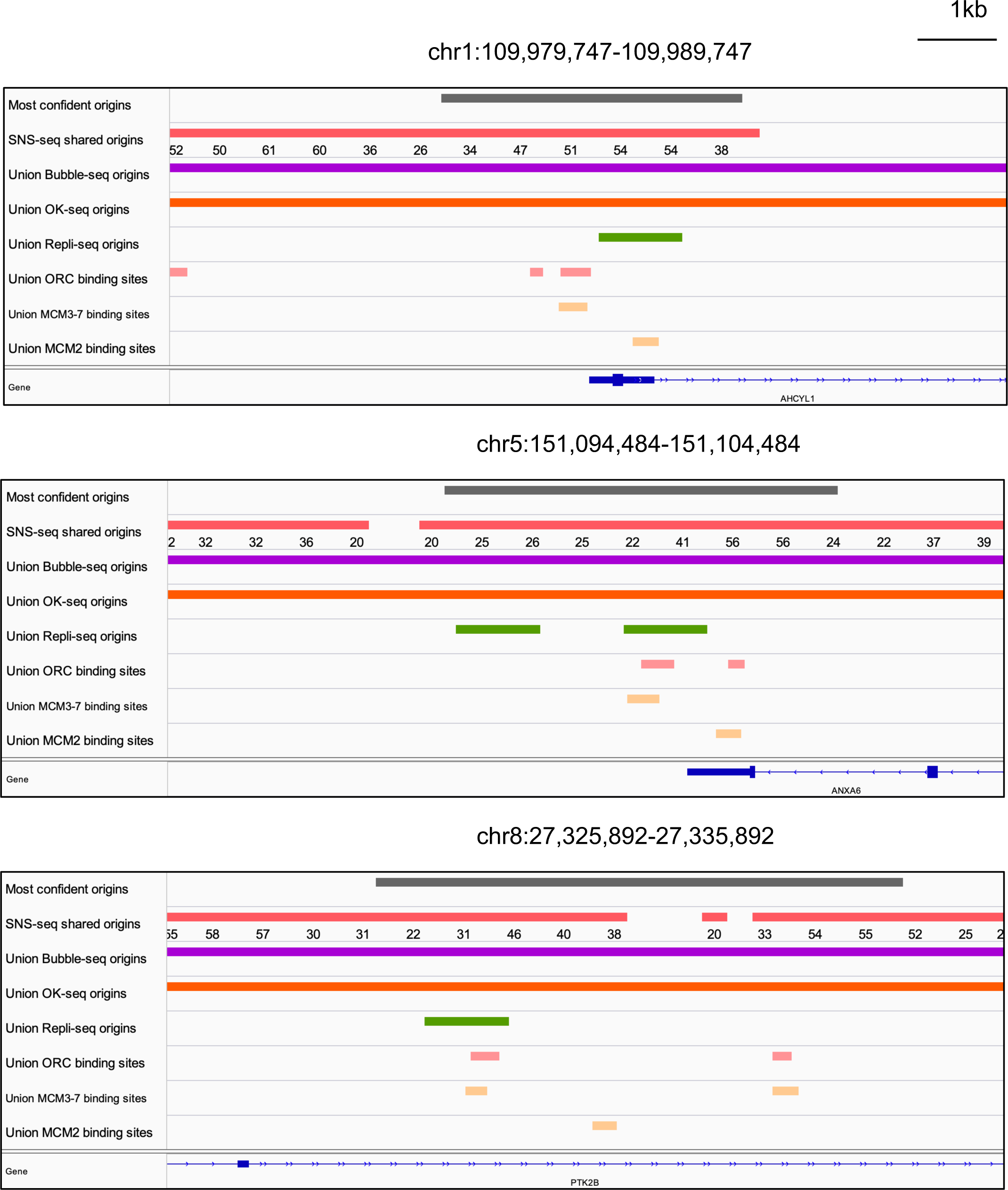
Genome browser screenshot for three of the 74 origins from Fig. 5h. The numbers below the SNS-seq shared origins track is the occupancy score of the origins along the length of indicated track.

## Discussion

Through integrative analysis of publicly available 113 profiles of replication origins from multiple techniques, we identified ∼7.5 million union origins in the human genome, of which only 20,250 shared origins are reproducibly identified by at least 20 SNS-seq datasets and confirmed by each of three other techniques. The following conclusions were reached: the shared (highly reproducible) origins are only a small subset of all origins identified and are in epigenetically active chromatin with a strong preference for promoters. However, G-quadruplexes, CTCF binding sites and ORC binding sites are also enriched near promoters, so that although these genomic features are enriched near the shared origins, it is not clear whether these genomic features actually specify origins, or whether they are simply enriched near origins because origins are near promoters. Finally, our results suggest that the co-localization of the shared origins with ORC or MCM2-7 binding sites is very low, much lower than in yeast, and so (a) the currently known binding sites of these proteins should not be used as surrogate markers of origins in human cells and (b) there is a great need to improve our methods of identifying isoforms of these proteins that are strictly functional for origin firing. The current models of origin firing require ORC to bind near origins, to load MCM2-7 nearby, and the latter to be converted to active helicase to fire origins. To rigorously test this model, we need examples of reproducible origins where these conditions are met, and our study helps identify 74 such sites where such experimental verification can be carried out.

Careful analysis shows that individual methods produce origin datasets that are more correlated with each other than across methods. The extensive variation of origins between datasets is likely because of background noise in all current techniques, and because there is extreme stochasticity of origin selection during DNA replication in individual cells and cell cycles in the same population of cells from a human cell line in culture. There are also differences that are created due to differences in cell lineage. Despite this, we could cut through those differences to focus on the small subset of shared origins that seem to be active in multiple cell lineages.

The shared origins are enriched with active histone marks and enriched in early replicating parts of the genome that are overwhelmingly in an active epigenetic environment. H3K4me3 has been reported to be enriched at replication origins (Picard et al. 2014; Cayrou et al. 2015), and it has also been reported that demethylation of H3K4me3 by KDM5C/JARID1C promotes origin active (Rondinelli et al. 2015). H3ac/H4ac are also reported to be enriched at replication origins (Cadoret et al. 2008; Sequeira-Mendes et al. 2009) and this is regulated by various histone acetyltransferases (HATs) and histone deacetylases (HDACS) (Goren et al. 2008; Wang et al. 2009). Interestingly, H3K27me3 has also been reported to be enriched at replication origins (Picard et al. 2014; Cayrou et al. 2015). The higher enrichment of *all* activating histone modifications relative to the repressive histone modifications (including H3K27me3) (Fig. 2j) rather than individual types of modification, suggest that the shared, highly reproducible origins, are preferentially in epigenetically active euchromatin, this aligns well with the general enrichment of shared origins with TSSs and TF hotspots, which are concentrated in gene-dense, epigenetically open parts of the chromosomes. This also aligns well with the fact that shared origins were not enriched for H4K20me3 and H3K9me3. H4K20me3 has been reported to be enriched near late-firing, heterochromatic initiation zones (Kirstein et al. 2021) and H3K9me3 enriched near late replicating origins (Wang et al. 2017), while the shared origins are biased towards those that are in euchromatin. That origins, like Transcription start sites, prefer areas of the chromatin marked by activating epigenetic marks has been reviewed in 2016 (Prioleau and MacAlpine 2016). We arrive at the same conclusion even though we worked with origins identified after this review, presumably with significant improvements in method and even though we focused on the most reproducible origins.

We characterized the genomic features of origins and found the shared origins are more co-localized with TSS. Table 1 shows the summary of the overlap of union origins, shared origins, shared origins overlapping ORC binding sites with various genomic features and compares this with the overlap of TSS with the same genomic features. Overall, these results suggest that as we proceed from all origins, to shared origins to highest confidence origins that are proximate to ORC binding sites, we see increasing overlap with promoters and early replicating, transcriptionally active, epigenetically open parts of the genome and that features thought to be important for origin selection (like G-quadruplexes, CTCF binding sites, active chromatin, ORC binding sites) could simply co-occur with origins because of their enrichment near promoters (TSS).

**Table 1.**
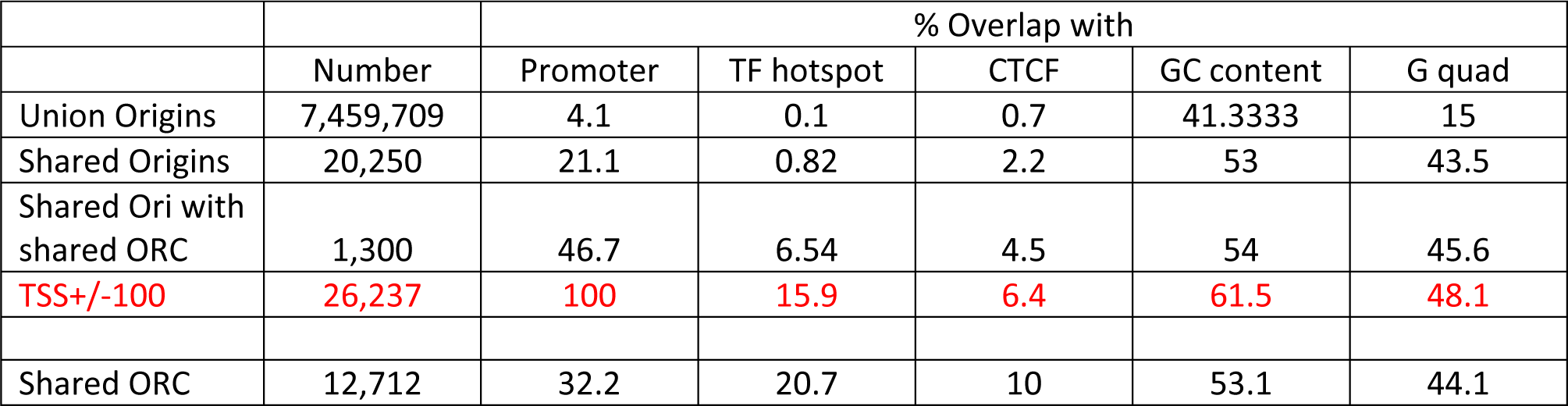
Overlap of origins, TSS and ORC binding sites with indicated features.

To rigorously test whether the overlaps of shared origins that we observe with various genomic features are significantly above background, we performed a permutation test that evaluates whether the observed overlap is significantly above the mean expected overlap when the experimental dataset is randomized 1,000 times (Table 2). The overlap of *shared origins* with promoters (TSS±4Kb), G-quadruplexes, R-loops, and CTCF binding sites are all significantly above background. However, the overlap of *promoters* with G-quadruplexes, R loops, and CTCF binding sites are also all significantly above background, though the enrichment of G-quadruplexes is minimal. Thus the genomic features characterized are all significantly enriched in both origins and promoters, and may help to specify both, either independently or co-dependently.

**Table 2.**
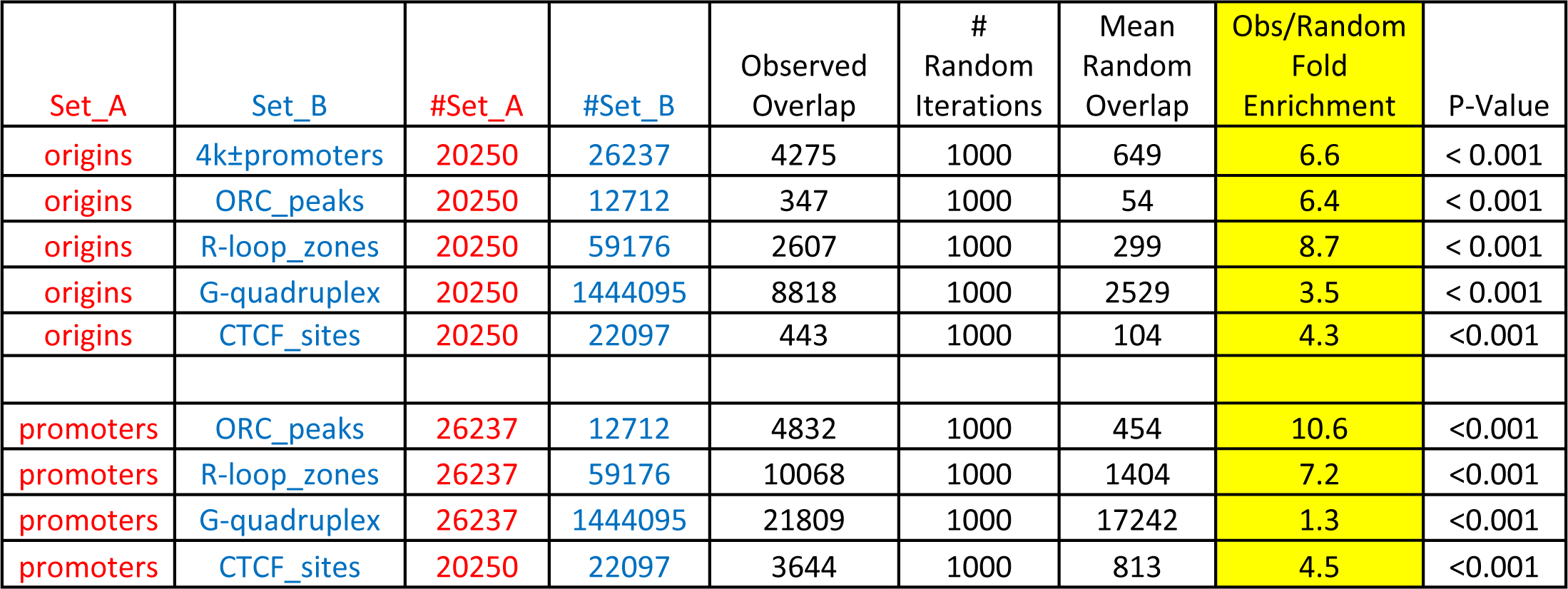
Permutation test of overlap of shared origins or promoters (TSS) with region around promoters, Shared ORC peaks (in > 2 datasets), R-loops, G-quadruplexes and CTCF binding sites. Fold enrichment of observed overlap relative to the mean overlap seen with 1000 randomizations of set A is indicated together with the p-value of the enrichment.

| Set_A | Set_B | #Set_A | #Set_B | Observed Overlap | # Random Iterations | Mean Random Overlap | Obs/Random Fold Enrichment | P-Value |
| --- | --- | --- | --- | --- | --- | --- | --- | --- |
| origins | 4k±promoters | 20250 | 26237 | 4275 | 1000 | 649 | 6.6 | < 0.001 |
| origins | ORC_peaks | 20250 | 12712 | 347 | 1000 | 54 | 6.4 | < 0.001 |
| origins | R-loop_zones | 20250 | 59176 | 2607 | 1000 | 299 | 8.7 | < 0.001 |
| origins | G-quadruplex | 20250 | 1444095 | 8818 | 1000 | 2529 | 3.5 | < 0.001 |
| origins | CTCF_sites | 20250 | 22097 | 443 | 1000 | 104 | 4.3 | <0.001 |
| promoters | ORC_peaks | 26237 | 12712 | 4832 | 1000 | 454 | 10.6 | <0.001 |
| promoters | R-loop_zones | 26237 | 59176 | 10068 | 1000 | 1404 | 7.2 | <0.001 |
| promoters | G-quadruplex | 26237 | 1444095 | 21809 | 1000 | 17242 | 1.3 | <0.001 |
| promoters | CTCF_sites | 20250 | 22097 | 3644 | 1000 | 813 | 4.5 | <0.001 |

G-quadruplexes are of particular interest because they have been experimentally shown to dictate origin-specification (Prorok et al. 2019; Valton et al. 2014). Although there is a higher overlap of shared origins than union origins with G-quadruplexes (43.5% vs. 15%) this overlap is not higher than that between promoters and G-quadruplexes (48.1%) (Fig. 2g). This may suggest that the overlap between origins and G-quadruplexes may be secondary to the overlap between origins and promoters. However, the overlap of shared origins with G-quadruplexes is 3.5-fold above random while that of promoters with G quadruplexes is only 1.3-fold above random, which would be consistent with the idea that G-quadruplexes have a role in specifying origins and the overlap is not simply due to the proximity of origins with promoters.

Similarly, CTCF binding sites have been proposed to contribute to origin-specification (Emerson et al. 2022). Here again, the 2.2% of shared origins that overlap with constitutive CTCF binding sites is less than the 6.4% of promoters that overlap with CTCF binding sites (Fig. 2e). Here, though, the permutation test in Table 2 reveals that the enrichment of origins near CTCF binding sites (4.3-fold above random background) is comparable to the enrichment of promoters near the same sites (4.5-fold). This may, thus, suggest that the overlap of CTCF binding sites with origins is secondary to the overlap of origins with promoters.

Since ORC is an early factor for initiating DNA replication, shared human origins should be proximate to the reproducible ORC binding sites. The vast majority of ORC binding sites are not proximate to the shared origins and conversely, only about 6.4% of the shared origins are proximate (within 1 kb) to the reproducible (shared) ORC binding sites (Fig. 3f). Even with the most relaxed criteria nearly 85% of the most reproducible origins in human cells are not proximate to any ORC binding site (union, Fig. 3f). This low level of proximity contrasts with the ∼100% of yeast origins that are within 1 kb of a yeast ORC binding site. Even when we examined cell line-specific origins with the ORC ChIP-seq datasets from the same cell lines, the co-localization between origins and ORC binding sites remained poor (Fig. 3-figure supplement 1f). Another study also noted that only 13% of SNS-seq origins in K562 cells are near ORC2 binding sites (Miotto et al. 2016), but the authors suggested that this could occur if many SNS-seq sites are not real origins but arose from DNA breaks. A vast contamination of SNS-seq origins by sites of DNA breaks is discounted in our analysis because of the reproducibility of the shared origins in multiple labs, multiple lineages and multiple different techniques.

Instead of an unbiased determination of origins mapped by multiple groups to create a small set of reproducible origins, we could also empirically select a few well-curated origin datasets. Towards that end we used the Core origins identified by lambda exonuclease SNS-seq (Akerman et al. 2020) and the Ini-seq2 origins identified by labeling nuclei *in vitro* to identify the earliest replicated parts of the genome (Guilbaud et al. 2022). The shared origins were only a small subset of these origin datasets (Fig. 5-figure supplement 3a and 3c). The percentage of these origins that were near ORC was still far less than what we observed in yeast: 13.7% for core origins and 30.4% for Ini-seq2 origins (Fig. 5-figure supplement 3b and 3d). The higher overlap with Ini-seq2 origins could be because *in vitro* initiation of replication in isolated nuclei in Ini-seq preferentially identifies origins in early replicating, euchromatic parts of the genome, areas that have already been reported to be more enriched for ORC2 binding sites (Miotto et al. 2016).

There could be other explanations for the poor overlap of shared origins with ORC binding sites. We tested whether increasing the stringency of reproduction in SNS-seq data will produce more reproducible origins that are better co-localized with ORC binding sites. However, when we reduced the shared origins based on their reproduction by 30, 40, or 50 (out of 66) SNS-seq datasets, we did not see a marked improvement of their co-localization with currently known ORC binding sites (Fig. 5-figure supplement 4a and 4b) and MCM binding sites (Fig. 5-figure supplement 4c and 4d).

Finally, the permutation test in Table 2 demonstrates that the co-localization of origins with ORC binding sites was 6.5-fold (p<0.001) greater than random expectation but this was less than the co-localization of promoters with ORC binding sites, 10.6-fold greater than random (p<0.001). Thus, the co-localization data cannot conclusively say whether ORC binding near an origin is because ORC specifies origins, or whether this is secondary to origins being co-localized near promoters. Note, that the co-localization of ORC1 binding sites and origins in HeLa cells with highly active TSS has been noted in the past (Dellino et al. 2013). In human lymphoblast Raji cells ORC was enriched in promoters, and MCM depleted from gene bodies (Kirstein et al. 2021). ORC1 has also been reported to bind to RNA near promoters and stimulate replication from such sites (Mas et al. 2023). Thus our observation that ORC is highly enriched near promoters is expected, the main advance being that the random permutation test suggests that co-localization of TSS with ORC is more than the co-localization of shared origins with ORC. The question arises, are our results contradicting past studies. To the best of our knowledge, no one has taken such a comprehensive and quantitative approach to assess the proximity of origins with ORC binding sites genome-wide. We determined the percentage of origins that are within 1 kb of ORC-binding sites genome-wide and did a random permutation test to test whether the observed overlap is greater than random expectation. This told us that only a small percentage of the origins are within 1 kb of currently known ORC binding sites on a genome-wide scale, although selected browser shots as in Fig. 6 might suggest otherwise. Furthermore, even though Origins were enriched near ORC binding sites relative to random expectation this enrichment is not more than that of TSS near ORC binding sites.

Taken together, these results suggest three possibilities: (a) the ORC ChIP-seq or origin determination datasets in humans are very noisy and that the ChIP-seq data in particular fail to identify functional ORC binding sites; (b) that the MCM2-7 loaded at the ORC binding sites move very far to initiate origins, farther than 1 kb from the ORC binding sites; or (c) that there are as yet unexplored mechanisms by which many of the most reproducible origins are specified, mechanisms that might include ORC-independent modes of origin specification, similar to the DnaA independent modes of origin specification in bacteria that use R-loops or DNA breaks to initiate replication (Itoh and Tomizawa 1980; Leela et al. 2021; Goswami and Gowrishankar 2022).

There is significant evidence for possibility (b) above. MCM2-7 is either loaded far from ORC binding sites or move significant distances after loading in Xenopus egg extracts (Harvey and Newport 2003). In yeasts there is evidence that MCM2-7 are pushed far away from the bodies of transcriptionally active genes by the RNA polymerase II and initiate replication at sites distant from where they are loaded on chromatin by ORC (Gros et al. 2015). However, it is worth noting that despite this nearly 100% of the origins in yeast are within 1 kb of ORC binding sites (Fig. 3f). Similar observations have been made in Drosophila, where cyclin E/cdk2 kinase activity promotes the loading of vast excess of MCM2-7 on chromatin relative to ORC, and the MCM2-7 complexes move away from their loading sites due to the activity of the transcriptional apparatus (Powell et al. 2015).

Since most models of replication initiation propose that a stably bound MCM2-7 complex is converted to an active CMG helicase at the time of origin firing, we hoped that even if ORC binding sites are not necessarily close to the shared origins, MCM2-7 binding sites will be more proximate to the shared origins in human cells. However, only 4.5% of shared origins are near any MCM2-7 binding site in human cells and again this is in contrast to the 96.5% of origins in yeast being near any MCM2-7 binding sites (Fig. 5g). Even if we focus on the limited data from the same cell line (K562 or HCT116) only 3.8% (union origins) or 5.5% shared origins overlap with union MCM3-7 binding sites in K562 cells and 19.5% of union origins overlap with any MCM2 binding sites in HCT116 cells (Fig. 3-figure supplement 1b, 1c and 1e). As with ORC, we also did the analysis with two selected origin datasets, the core origins (Akerman et al. 2020) and the Ini-seq2 origins (Guilbaud et al. 2022) and found that still a small percentage of these origins (6.6% and 10.7%, respectively) were near any MCM2-7 binding sites. Finally, a recent paper reported the binding sites of phosphorylated isoforms of MCM2 (Thakur et al. 2022), and since phosphorylation of MCM2 is a prerequisite for MCM2-7 being activated as a helicase, we asked whether any of the 24hosphor-MCM2 binding sites showed better co-localization with the shared origins (Fig. 5-figure supplement 2). Phospho-S108 MCM2 showed the best result among the phospho-isoforms tested, with 22% of the shared origins being proximate to the Phospho-S108 MCM2 binding sites. Thus, phosphorylation on S108 of MCM2 may indeed mark MCM2-7 complexes that become active helicases which fire origins, but even then the co-localization was seen in a disappointingly low percentage of shared origins.

MCM2-7 ChIP-seq data is biased by a lot of noise as human MCM2-7 slides around after initial loading and due to contamination from the actively replicating CMG helicase that moves all over the genome (and pauses at many sites) during S phase, but this should increase the number of MCM2-7 binding sites and not decrease the percentage of origins that are near MCM2-7 sites. The better co-localization of shared origins with phospho-S108 MCM2 suggests that the ChIP methods can be improved to identify a small but critical pool of MCM2-7 that is engaged with the DNA stably in a way that permits origin specification. Alternatively, mapping of the sites bound to the active CMG helicase (MCM2-7, CDC45 and GINS), before the active helicase moves too far from the initiation site, may better enrich for sites that are destined to be origins.

A byproduct of our analysis of ORC ChIP-seq data, was the discovery that ORC1 and ORC2 are not strictly co-located as expected for the subunits of one ORC complex. Only 1,363 (21%) of the 6,501 ORC1 peaks overlapped with the 29,930 ORC2 peaks (Fig. 5-figure supplement 1a), and the vast majority (68%) of ORC1 peaks are far away (≥2kb) from the closest ORC2 peaks (Fig. 5-figure supplement 1b). As a positive control, we checked other proteins that form well-established complexes like SMARCA4 and ARID1A in SWI/SNF complex, SMC1A and SMS63 in 25ohesion complex, EZH2 and SUZ12 in PRC2 complex, and found that they show a high proportion of overlap between their peaks as expected (Fig. 5-figure supplement 1a). A possible explanation of the low overlapping rate between ORC1 and ORC2 peaks could be that the datasets are from different cell types (Fig. 5-figure supplement 1c). However, even in ChIP-seq peaks from different cell types, we found continued high overlap between SMC1A and SMC3 peaks as would be expected for complexes that do not have high inter-cell-line variability in binding sites. In contrast, there was a low overlap between EZH2 and SUZ12 peaks taken from different cell-lines suggesting that for other complexes there is significant inter-cell-line variability in binding sites (Fig. 5-figure supplement 1d). Thus, the lack of overlap between the ORC1 and ORC2 association sites on chromatin could be explained by inter-cell-line variation of binding sites for ORC (like the PRC2 complex) or could be explained by the ORC subunits binding to chromatin independent of one another. We prefer the latter explanation because of our experimental result that a substantial fraction of ORC2, 3, 4, 5 and 6 bind to chromatin in the absence of ORC1, and ORC2, 3 and 6 bind to chromatin normally in the absence of ORC5 (Park and Asano 2008; Shibata et al. 2016; Okano-Uchida et al. 2018; Shibata and Dutta 2020). We and others have also noted that human ORC2, 3, 4 and 5 form a stable subcomplex, with much looser association of the subcomplex with ORC1 and ORC6 (Dhar et al. 2001). Note that lineage-specific variation of where ORC may bind to the chromatin or the binding of ORC subcomplexes to chromatin may explain why so few of the ORC binding sites overlap with the shared origins but does not explain why only 6.4% of the shared origins (identified reproducibly in different cell lines) is proximate to any ORC binding sites.

Our analysis provides a characterization of origins in the human genome using multi-source data in the public domain. The sequencing-based profiles from different techniques have different biases in detecting DNA replication origins and show different genomic features. As one adds criteria to identify the most reproducible experimentally determined origins that are close to ORC binding sites, the overlap with TSS, TF-hotspots, constitutive CTCF binding sites and G-quadruplexes increases (Table 1), but in general the overlap with the latter three genomic features does not exceed that of the same features with TSS. Thus, although the overlaps are statistically significant, this correlation analysis cannot determine whether the same genomic features *independently* specify origins of replication and promoters of transcription. This is a question that should be asked experimentally and the 74 experimentally reproduced origin datasets with proximity to ORC and MCM2-7 ChIP-seq sites provides a starting list for such experimental queries. Finally, although our datasets for origins were large, as with all integrative data analyses we are limited by the quality of the data, which can be improved as experimental techniques for both origin identification and ORC or MCM2-7 binding site determination continue to improve.

## Methods

### Data processing

We collected all publicly available human sequencing datasets of replication origin profiling and ORC ChIP-seq from GEO database (Barrett et al. 2013). Data details of the collected datasets can be found in Supplementary File 1. Raw sequence data in fastq format were downloaded and processed as follows: FastQC (Andrews S 2010) (v0.11.5) was used for quality control and sequence data were then mapped to human genome (hg38) using bowtie2 (Langdon 2015) (v2.2.9). Sam files were converted into bam files using samtools (Li et al. 2009) (v1.12) and only high-quality reads (q-score ≥30) were retained for subsequent analyses. Peak calling for ORC ChIP-seq data was performed with MACS2 callpeak function (--nomodel –extsize 150 -g hs -B – SPMR -q 0.05 –keep-dup 1) (Zhang et al. 2008) (v2.1.4). Samples with more than 1000 peaks were kept as high-quality samples. SNS-seq, Bubble-seq, and Rerep-seq peak calling was performed with SICER (Zang et al. 2009) with different parameters based on the different resolution of technique: SNS-seq (W200 G600), Bubble-seq (W5000 G15000), Rerep-seq (W200 G600). The OK-seq defined Izs is generated from previous published papers (Petryk et al. 2016), (Wu et al. 2018) and the coordinates information is provided by Drs. Chunlong Chen and Olivier Hyrien. Repli-seq defined origins is generated from UW Encode group and is accessible in GSE34399. To obtain union peaks, peaks from each technique and from all techniques were merged using Bedtools (Quinlan and Hall 2010) merge (v2.29.2). Union origins that are longer than 300bp were cut into separate origins of a maximum length of 300bp. At the end we obtained 7,459,709 union origins.

To compare the locations of origins between samples, we performed a principal component analysis (PCA), where the matrix (to which PCA is applied) is of size 7.5M (number of union origins) x 113 (samples), with a “1” in the (*I*,*j*) position if the *i*th origin was detected in the *j*th dataset. We also performed pairwise pearson correlation of the origins in all the datasets to determine the reproducibility of the results from a given technique and across techniques. The distance metric for heatmap is generated in R using function cor (with parameters: use = “pairwise.complete.obs”, method = “pearson”).

### Identification of shared origins

The origins detected by each sample are very different, which makes the identification of common origins difficult. SNS-seq origins have the highest resolution, and so we start with them. We identified 5,069,432 union SNS-seq origins using the same approach. We focus on the union SNS-seq origins that appear above a threshold number of SNS-seq origin datasets (high confidence SNS-seq origins) and then determine those that overlap with any initiation zone identified by each of the three different techniques to delineate a set of “shared origins”, that initiate replication all the time and can be detected by multiple approaches. Note that this approach does not lose the high resolution of the SNS-seq origins, it merely finds those SNS-seq origins that are most reproduced and detected by each of the other 3 techniques. Rerep-seq is not included in define shared origins because it is very different with other techniques in many ways (Fig. 1 b-h, Fig. 1-figure supplement 1g).

To identify the most commonly shared origins between SNS-seq samples, we use an occupancy score for each union SNS-seq origin to show how many samples identify that specific origin. A higher occupancy score means the origin is more commonly present in many samples (Fig. 2-figure supplement 1a-b). High confidence SNS-seq origins are those that occur in a sufficient number of SNS-seq samples at an FDR of >0.1. We used an exponential model for background (green line) and plotted the distribution of occupancy scores (blue dotted curve) for origins from all SNS-seq samples (Fig. 2a). The exponential model can be expressed as

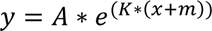

where x is the occupancy score, and y is the expected number of origins. Origins that are reproduced by multiple samples have higher occupancy scores than the background distribution. An empirical FDR of 0.1 was used to determine the cutoff of occupancy score so that the number of observed origins with occupancy score greater than the cutoff should be 10 times more than expected in the background model (Fang et al. 2020). The high-confidence SNS-seq origins were thus those with occupancy score ≥20 in all 66 SNS-seq samples. Detailed parameters can be found in Supplementary File 2. Origins mapping to ChrM are removed from SNS-seq shared origins to avoid the interference of mitochondria DNA.

Finally, the high-confidence SNS-seq origins defined by our model that overlap with any IZs from OK-seq, any IZs from Bubble-seq, and any origins defined by Repli-seq are called “shared origins” (Fig. 2b). A total of 20,250 shared origins were identified.

K562 shared origins were identified as K562 SNS-seq origins that overlapped with any of the union OK-seq and Repli-seq peaks. HeLa shared origins were identified as SNS-seq origins present in all 3 SNS-seq samples in HeLa and overlapped with any of the union OK-seq and Repli-seq peaks.

### Origins, ORC and MCM ChIP-seq data in yeast

Yeast origins data were obtained from OriDB (Nieduszynski et al. 2007). 829 replication origins mapped to the yeast genome sacCer1 were converted to the sacCer3 genome using UCSC LiftOver (Hinrichs et al. 2006). 289 experimentally confirmed origins, considered as high-confidence origins, were used for co-localization analysis with ORC and MCM binding sites in yeast. ORC and MCM ChIP-seq datasets were collected from the public domain (Supplementary File 1) and processed using the same procedure as used for human data with reference genome version sacCer3. The shared MCM binding sites were defined as the MCM peaks that occur in all samples.

### Enrichment of histone marks at shared origins

Histone modification ChIP-seq peak files were downloaded from CistromeDB (Zheng et al. 2019). A total of 5,711 peak files, each with over 5,000 peaks, covering 29 distinct histone modifications in different human cell types were used to interrogate the association between histone marks and shared origins, using union origins as control. The enrichment analysis was applied for each peak file by comparing the peak number in each histone modification peak file overlapping with shared origins versus the overlapping with union origins. The Odds ratio obtained from two-tailed Fisher’s exact test was used as the enrichment score for each peak file (Fig. 2j). Odds ratio and pvalue are calculated using python package scipy. (stats.fisher_exact([[shared_ori_overlap_with_hm, shared_ori_not_overlap_with_hm], [all_ori_overlap_with_hm - shared_ori_overlap_with_hm, all_ori_not_overlap_with_hm - shared_ori_not_overlap_with_hm]]))

### Co-localization analysis

We use co-localization analysis to define shared origins and show the overlapping of CTCF peaks, TF hotspots, TSS regions, G quadraplex peaks, etc. with origins. The co-localization analysis was performed using Bedtools (Quinlan and Hall 2010) intersect -u. At least 1 base pair of intersection is required to be defined as overlapping.

### Distance to the closest feature

According to the fact that ORC usually binds to origins but the loaded MCM2-7 can shift before firing an origin, we defined ORC that binds near the origins +/- 1kb as ORC near origins. Bedtools closest -d -t “first” was used to find the closest peak/region and distance for a given region. Origins with a binding site of ORC no further than 1kb are selected as origins near ORC binding site.

### G4 sites

G4(G-quadraplex) predicted motif sites are obtained from a published G4 motif predictor: G4Hunter(Bedrat et al. 2016). The 1,444,095 G4 motif coordinates is provided by Laurent.

### Genomic annotation of promoter, exon, intron, and intergenic region coverage

The coordinates of TSS, exon, intron are from UCSC hg38 version(Navarro Gonzalez et al. 2021). Promoter regions are defined as TSS+/-1kb. Intergenic regions are other genome regions excluding promoter, exon, and intron regions. Genomic annotation for peaks is defined by overlapping with those 4 types of regions by at least 1 bp.

### Replication timing score

We used publicly available Repli-seq data in K562 cell line for different cell phase from ENCODE project(Navarro Gonzalez et al. 2021) to measure the replication timing of origins(Supplementary file 1). We used a previously defined weighted average score(Hansen et al. 2010) to combine signal from the six cell phases with the following formula: score=(0.917*G1b)+(0.750*S1)+(0.583*S2)+(0.417*S3)+(0.250*S4)+(0*G2). Higher values correspond to earlier replication.

### Permutation Test

The R (version 4.1.3) package named regioneR (version 1.24.0) was used to statistically evaluate the associations between region sets with minor modification. The following parameters were used for running regioneR. Number of iterations: 10,000. Evaluation function: numOverlaps. Randomization function: randomizeRegions. randomizeRegions: hg38. non.overlapping: TRUE. mc.set.seed: FALSE

## Acknowledgement

This work was supported by R01 CA60499 to AD and R35 GM133712 to CZ and K99 CA259526 to ZS. We thank all members of both the Zang and Dutta Labs for many helpful suggestions.

## Supplementary data

**Fig. 1-figure supplement 1.**
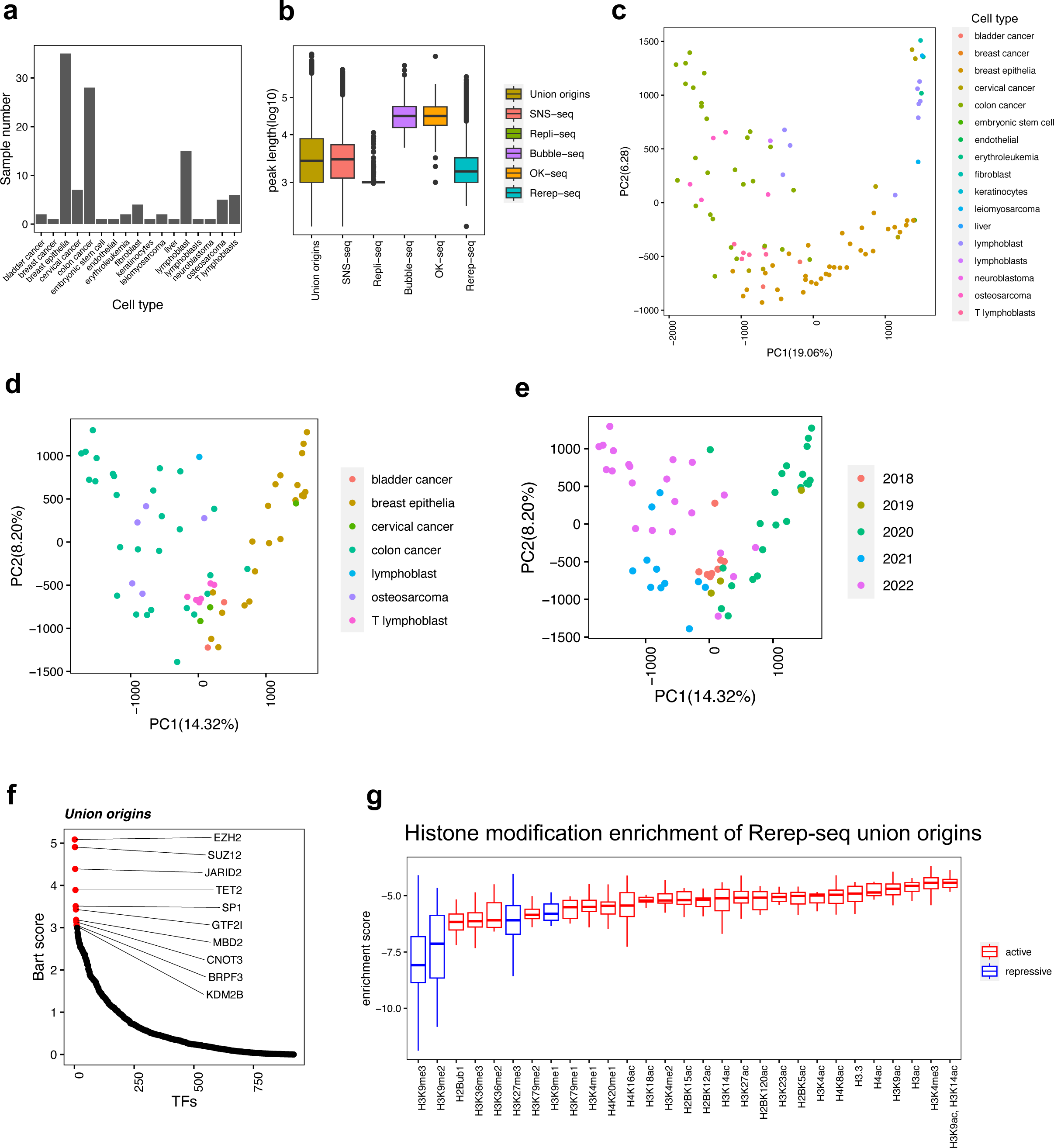
Distribution of origins defined by four types of techniques. (a) For each cell type, how many sample we have collected. (b) Distribution of peak length of origins from each technique. (c) PCA results of all samples, marked by cell types. (d) PCA results of SNS-seq samples, marked by cell types. € PCA results of SNS-seq samples, marked by year of the data uploaded. (f) BART2 results of union origins. (g) Enrichment of histone marks at re-replicated union origins using total union origins as control.

**Fig. 1-figure supplement 2.**
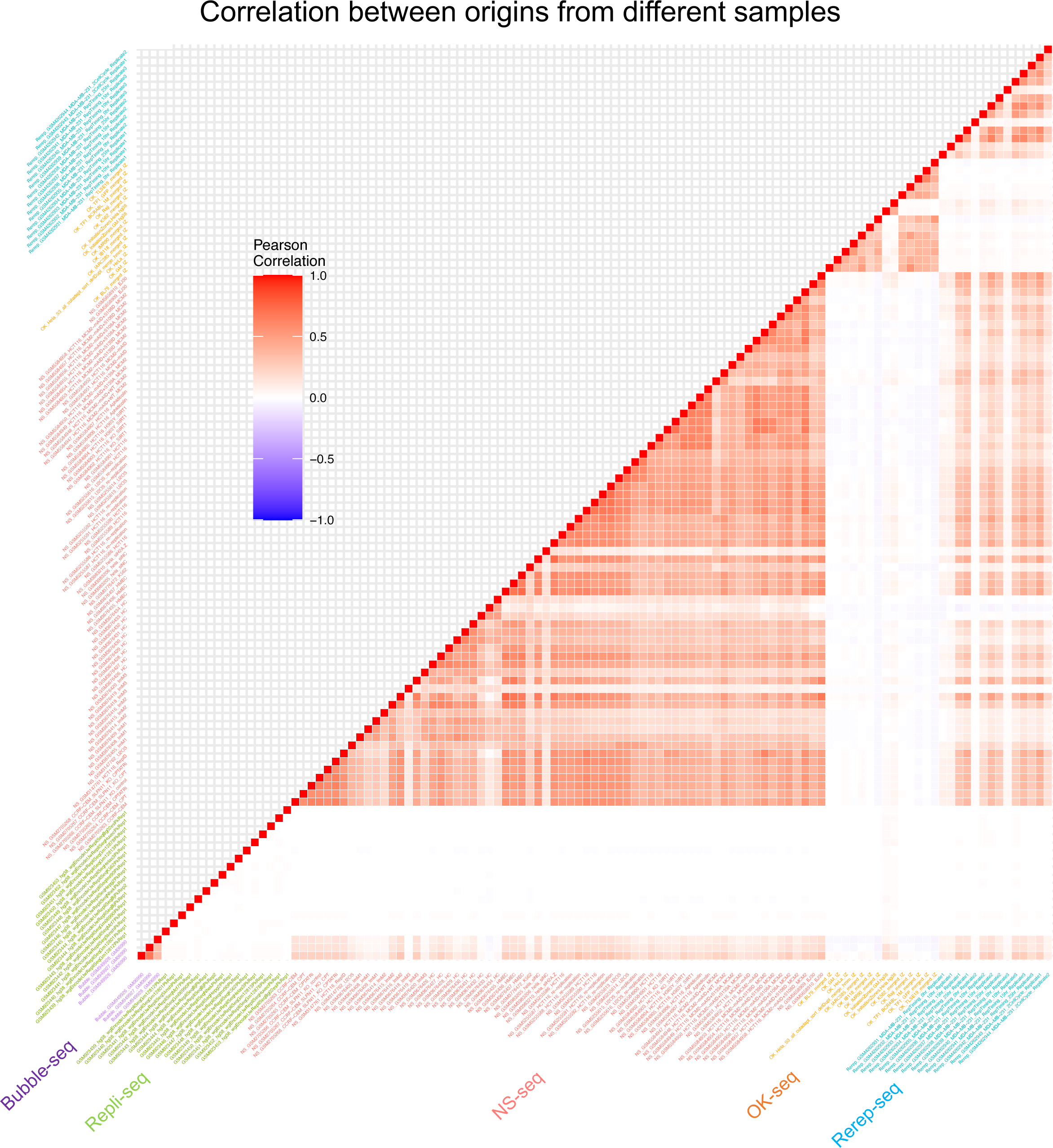
Correlation between origins from different samples. Pairwise correlation of samples from different techniques.

**Fig 2-figure supplement 1.**
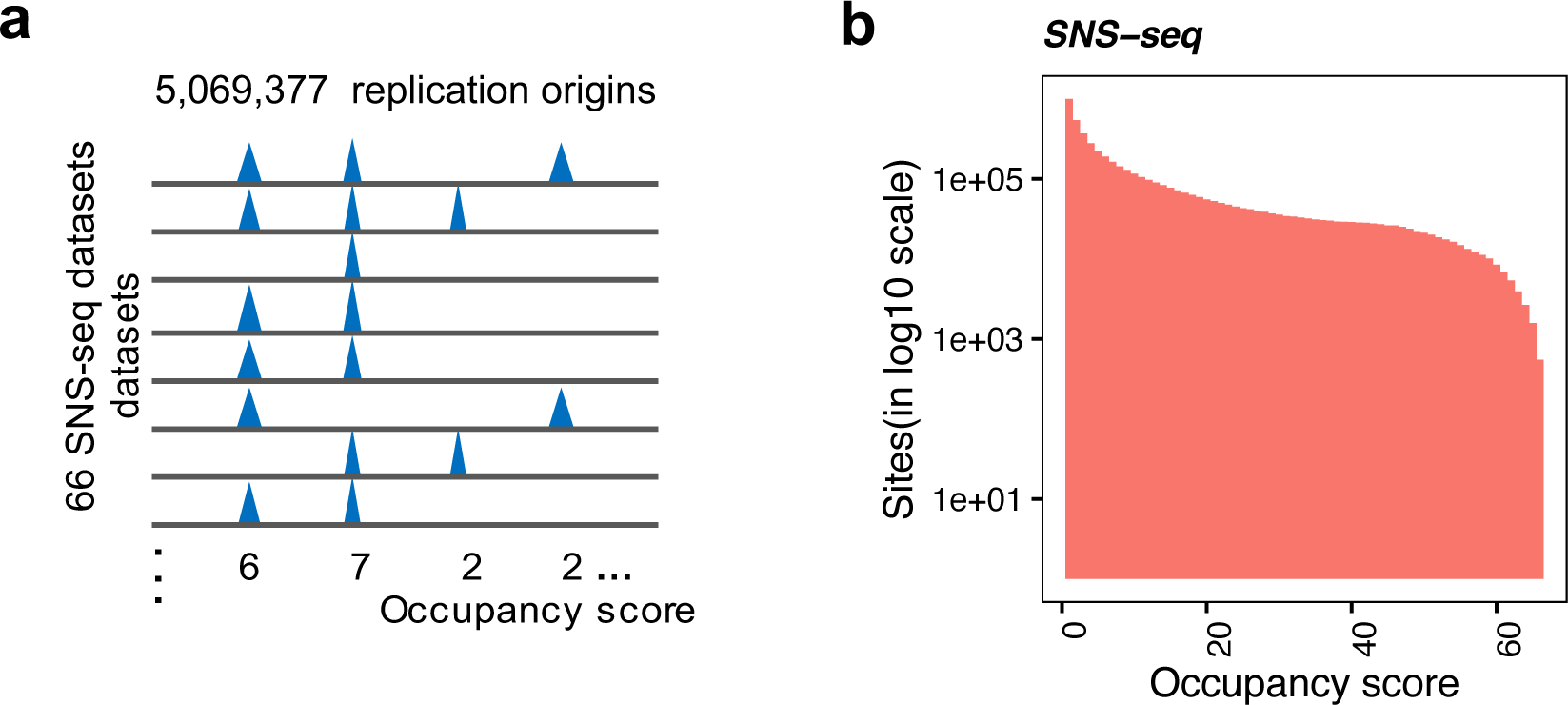
Background model for identification of shared origins. (a) Conceptual model of how occupancy score is defined to represent the number of samples that each origin occurs. (b) Distribution of occupancy score of SNS-seq union origins (300bp).

**Fig. 2-figure supplement 2.**
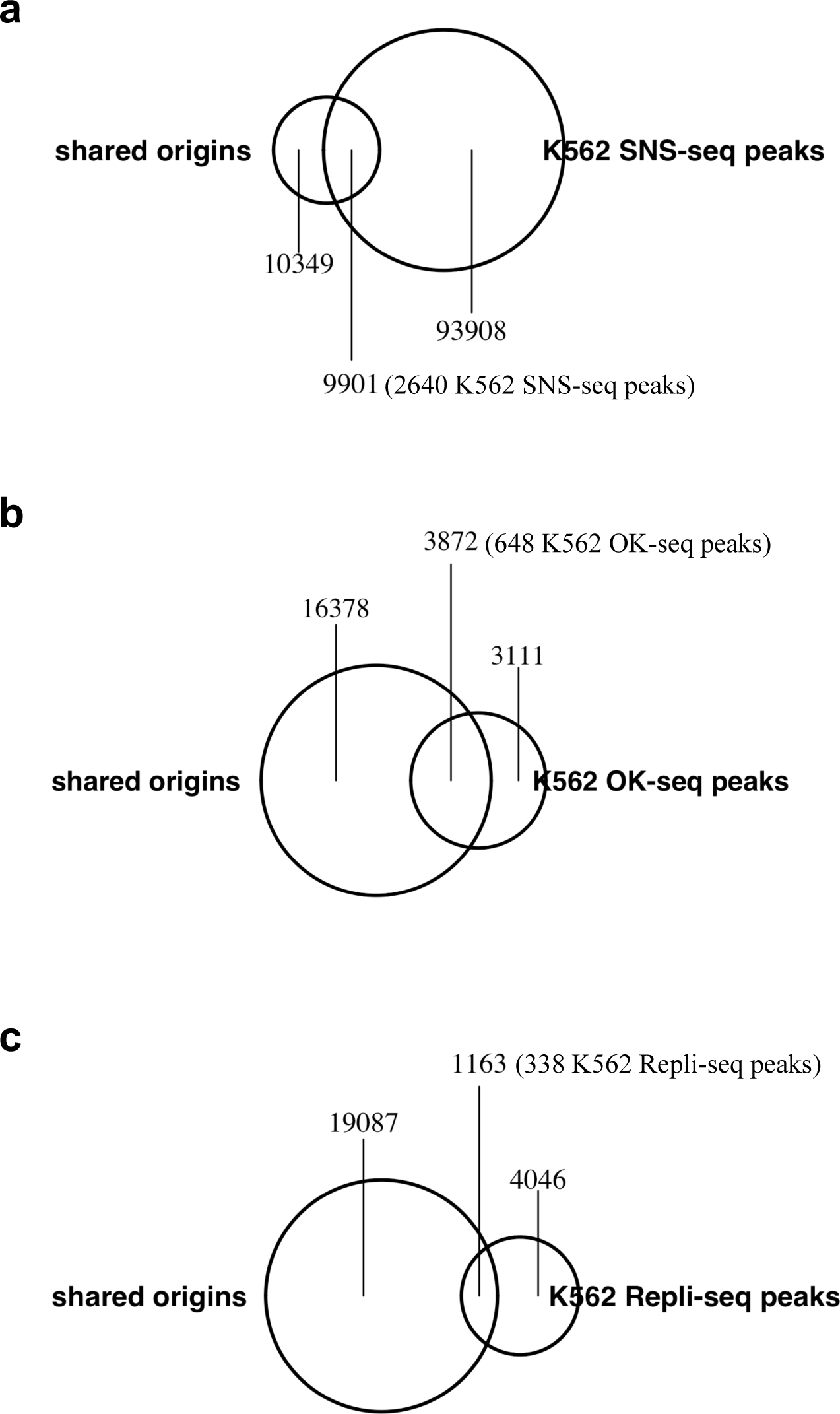
Origins defined by different techniques in K562 cell line and their overlap with shared origins. Shared origins are defined from all samples. The number of shared origins covered by each file is calculated and marked in the figure. Numbers in the parentheses are the numbers of peaks in the other dataset that overlap with the shared origins.

**Fig. 3-figure supplement 1.**
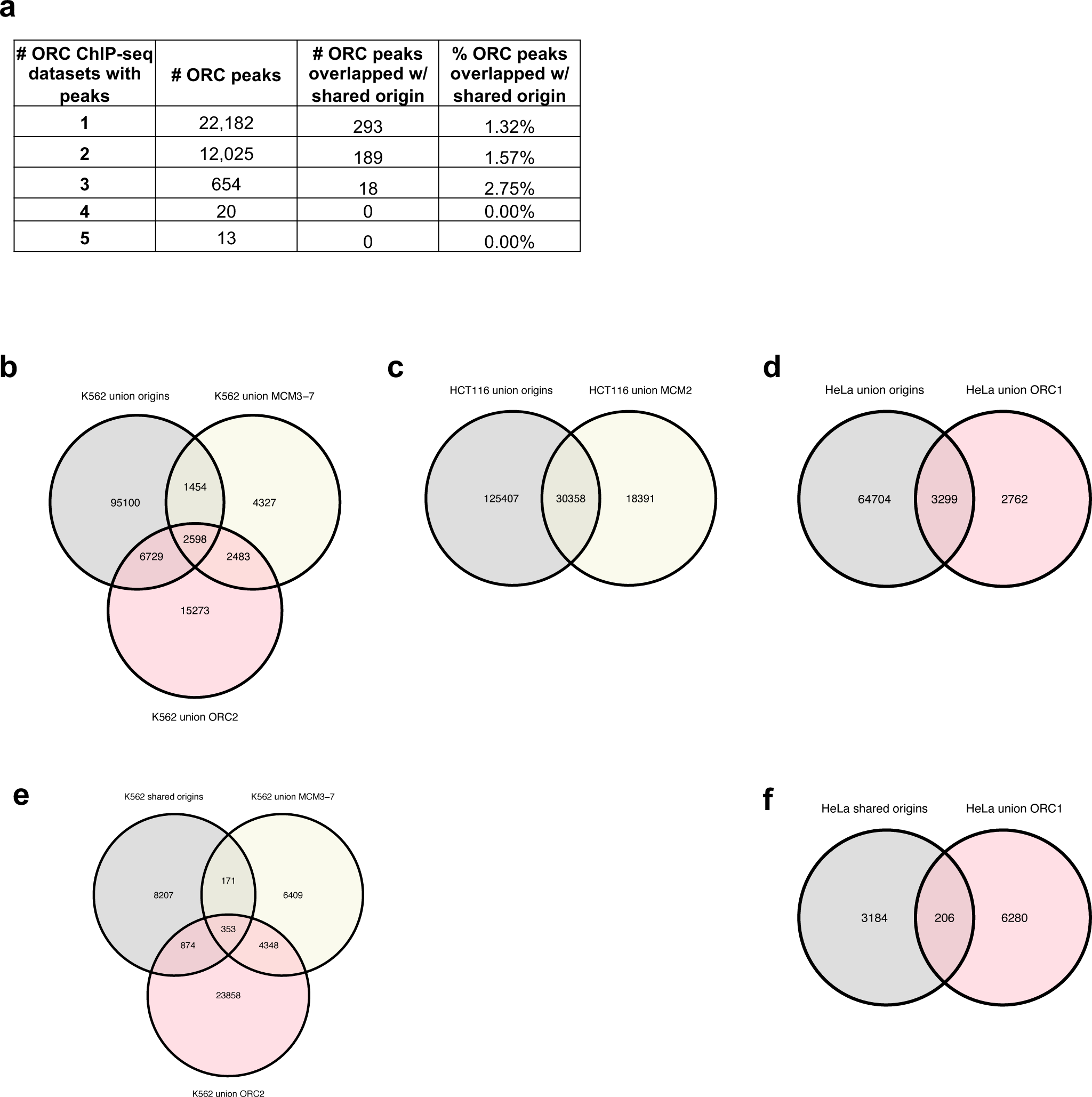
Analysis of overlap between shared ORC binding sites and origins. **(a)** 12,712 ORC binding sites in the human genome were shared by at least 2 ORC ChIP-seq datasets. The overlapping rates with shared origins is provided. **(b)** Overlapping of union origins, MCM3-7, and ORC2 in K562 cell line. **(c)** Overlapping of union origins and MCM2 in HCT116 cell line. **(d)** Overlapping of union origins and ORC1 in HeLa cell line. **€** Overlapping of shared origins seen in K562 cells with ORC and MCM binding sites in K562 cells. Shared origins seen in K562 cells are generated from SNS-seq files that overlapped with K562 Izs (defined by OK-seq and Repli-seq). **(f)** Overlapping of shared origins seen in HeLa cells with ORC binding sites in HeLa cells. Shared origins seen in HeLa cells are generated from 3 HeLa derived SNS-seq samples using the intersected peaks from: NS_GSM3983205_hela_siNC.bed, NS_GSM3983206_hela_siNC.bed, NS_GSM3983210_hela_siH2A.Z.bed.

**Fig. 5-figure supplement 1.**
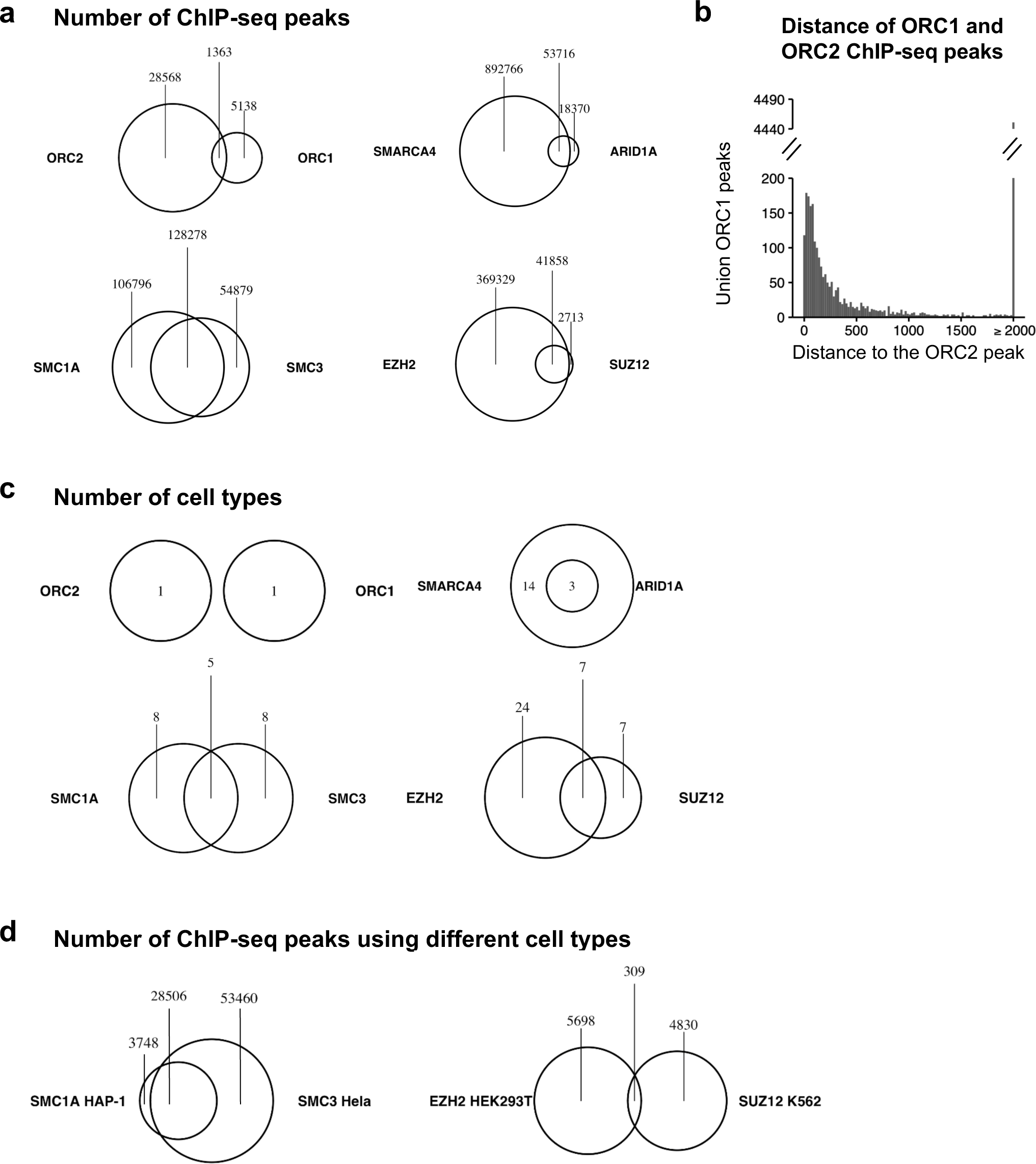
ORC subunits do not co-bind to DNA as much as expected. (a) Overlap of ChIP-seq peaks of different co-factors. (b) Distance distribution of ORC1 and ORC2 binding sites. (c) Number of shared cell types of ChIP-seq data used in (a). (d) Overlap of ChIP-seq peaks of co-factors from different cell types.

**Fig. 5-figure supplement 2.**
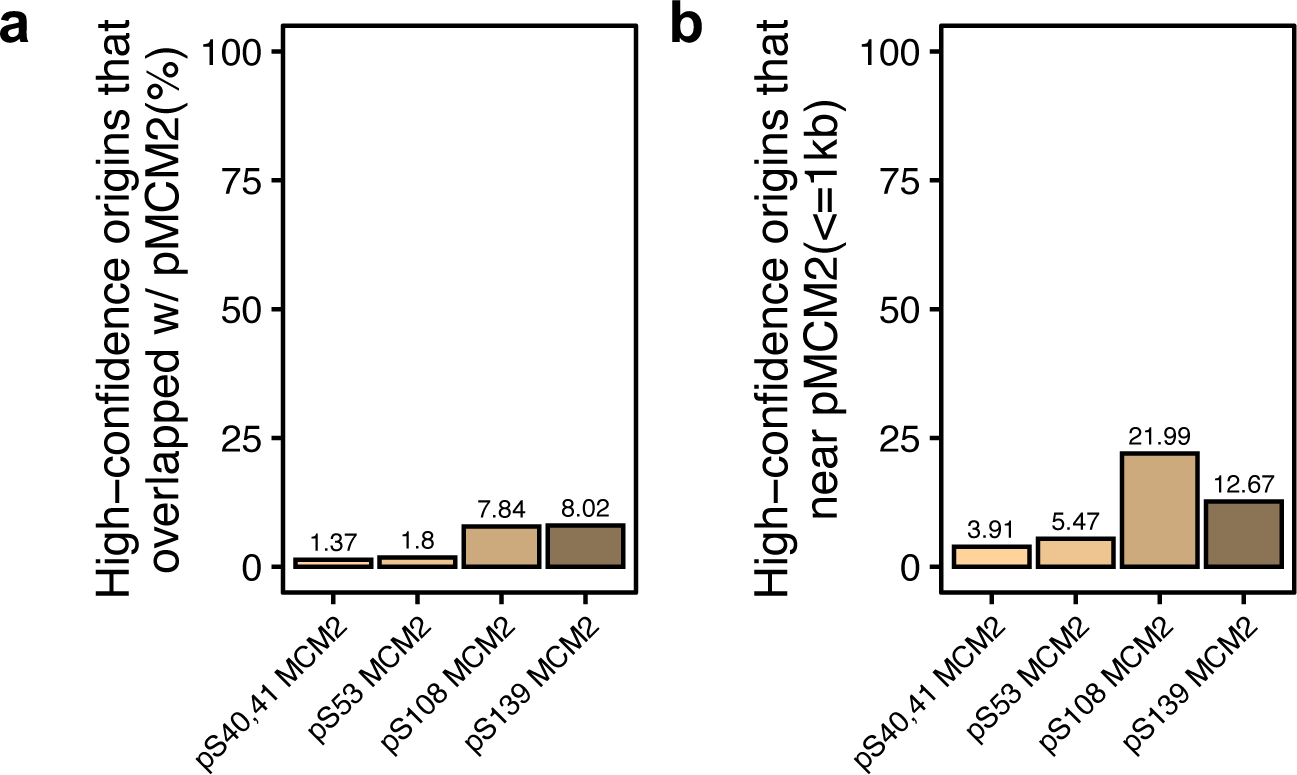
Shared origins overlap with phosphorylated MCM2. **(a)** Percentage of shared origins that overlap with phosphorylated MCM2 binding sites. **(b)** Percentage of shared origins that near phosphorylated MCM2 binding sites.

**Fig. 5-figure supplement 3.**
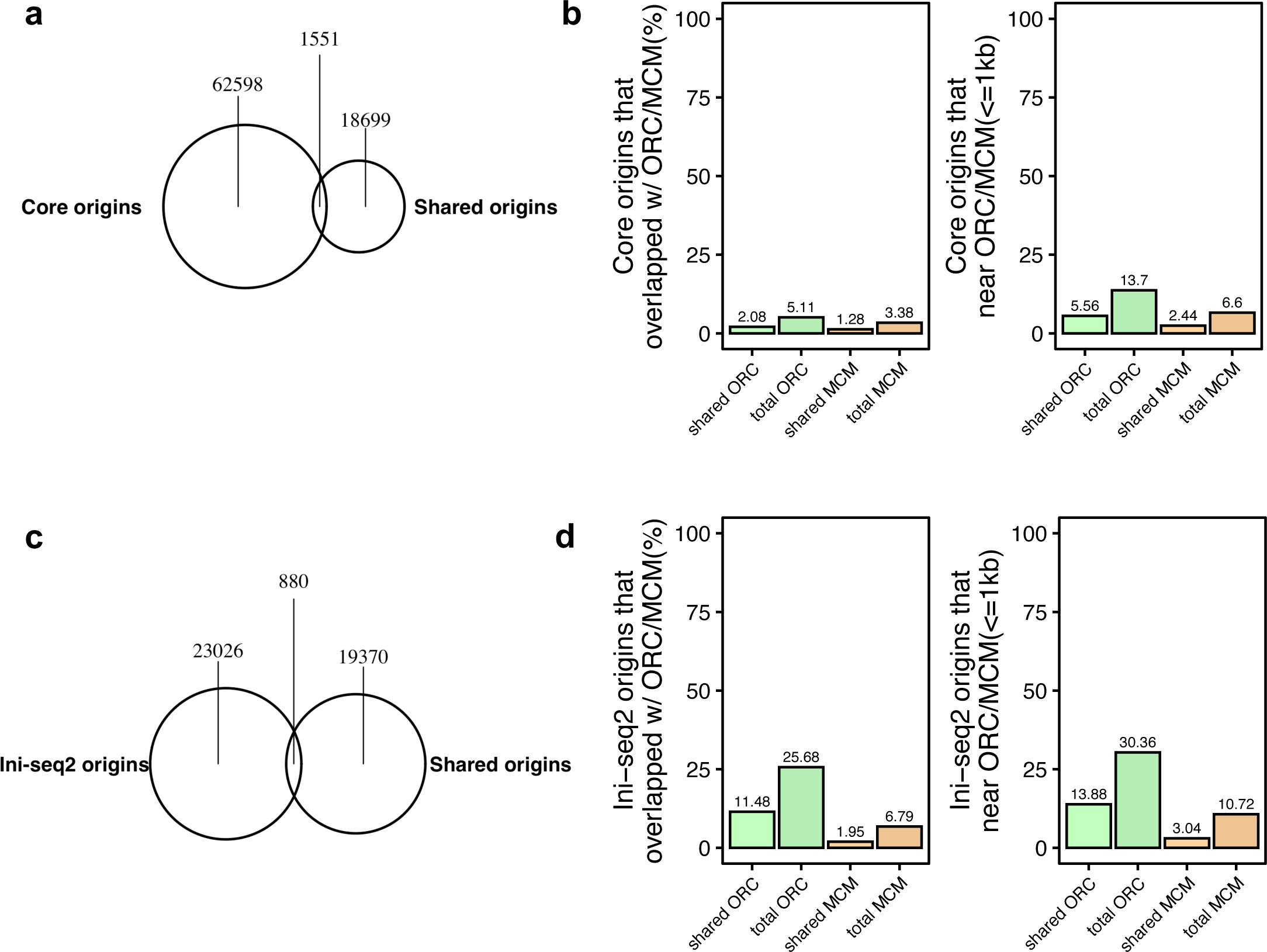
Analyses of a few selected origin sets suggested by reviewers. **(a)** Overlapping number of shared origins and core origins (Akerman et al., Nature Communications,2020). **(b)** Percentage of shared origins that overlap with phosphorylated MCM2 binding sites. **(c)** Overlapping number of shared origins and Ini-seq2 defined origins. **(d)** Percentage of shared origins that near phosphorylated MCM2 binding sites.

**Fig. 5-figure supplement 4.**
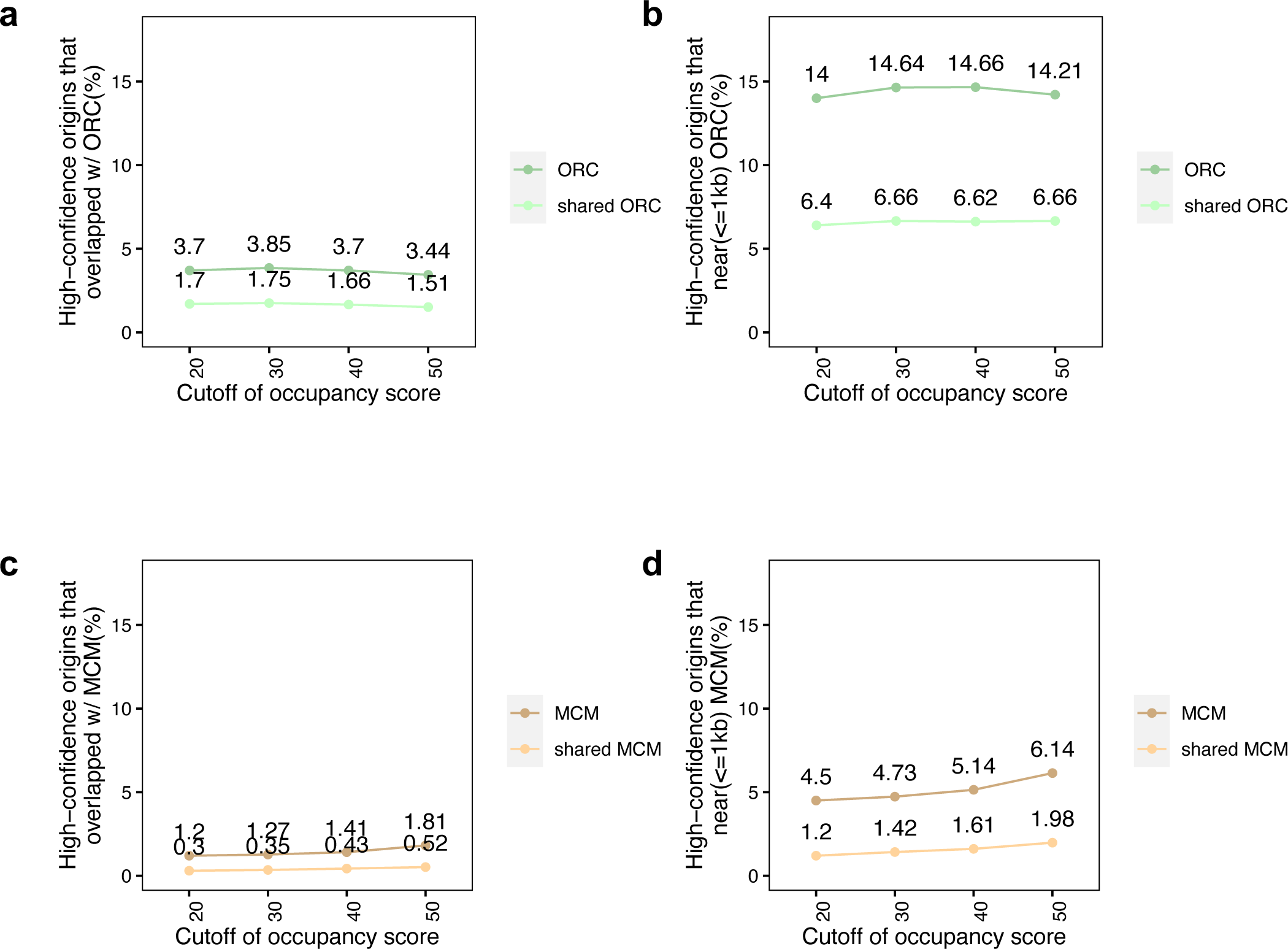
Selecting fewer but even more reproducible origins with more stringent cutoff to determine their overlap with ORC and MCM binding sites. **(a)** The percentage of high-confidence origins (defined by cutoff of occupancy score indicated on X-axis) that overlapped with union or shared ORC binding sites. **(b)** Similar to (a), except % origins that are near (< 1kb) ORC binding sites. **(c)** Similar to (a) except % origins that overlap with union or shared MCM binding sites. **(d)** Similar to (c), except % origins near (<1 kb) MCM binding sites.

**Supplementary File 1**, collected public data. The original reference for each dataset can be found on the GEO page for each dataset.

a, Metadata of collected public Origin data. b, Sample number of each cell type.

c, Metadata of collected public ORC ChIP-seq data. d, Metadata of collected public MCM ChIP-seq data.

e, Metadata of collected public ORC and MCM ChIP-seq data in yeast.

**Supplementary File 2,** parameters of model for SNS-seq Parameters of exponential model for SNS-seq origins.

**Supplementary File 3,** union and shared ORC binding sites

a, Union ORC ChIP-seq peaks, with coordinates and occupancy scores.

b, Shared ORC binding sites (defined as union ORC ChIP-seq peaks with an occupancy score >=2).

**Supplementary File 4,** highest confidence origins

a, Coordinates of union ORC ChIP-seq binding sites with shared origins in 1kb region.

b, Coordinates of shared origins with shared ORC binding sites in 1kb region.

c, Coordinates of shared origins with union ORC binding sites in 1kb region.

**Supplementary File 5,** 74 most confident origins

Coordinates of 74 origins that were reproducibly identified by multiple methods (shared origins, near shared ORC binding sites, overlapping with MCM3-7 binding sites and MCM2 binding sites).

